# Mapping histologic and functional maturation of human endocrine pancreas across early postnatal periods

**DOI:** 10.1101/2024.12.20.629754

**Authors:** Diane C. Saunders, Nathaniel Hart, Fong Cheng Pan, Conrad V. Reihsmann, Alexander L. Hopkirk, Nike Izmaylov, Shaojun Mei, Brandon A. Sherrod, Corey Davis, Jeff Duryea, Rachana Haliyur, Radhika Aramandla, Heather Durai, Greg Poffenberger, Alexandra Martin, Amanda L. Posgai, Irina Kusmartseva, Maria L. Beery, Mingder Yang, Hakmook Kang, Dale L. Greiner, Leonard D. Shultz, Jean-Philippe Cartailler, Kristie I. Aamodt, Rita Bottino, Mark A. Atkinson, Christopher V.E. Wright, Alvin C. Powers, Marcela Brissova

**Affiliations:** Division of Diabetes, Endocrinology and Metabolism, Department of Medicine, Vanderbilt University Medical Center, Nashville, Tennessee, 37232, USA; Department of Cell and Developmental Biology, Vanderbilt University, Nashville, Tennessee, 37232, USA; Department of Pathology, Immunology, and Laboratory Medicine, College of Medicine, University of Florida, Gainesville, Florida, 32610, USA; Department of Biostatistics, Vanderbilt University Medical Center, Nashville, TN 37232, USA; Department of Molecular Medicine, Diabetes Center of Excellence, University of Massachusetts Chan Medical School, Worcester, Massachusetts, 01655, USA; The Jackson Laboratory, Bar Harbor, Maine, 04609, USA; Vanderbilt Center for Stem Cell Biology, Vanderbilt University, Nashville, Tennessee, 37232, USA; Division of Pediatric Endocrinology, Department of Pediatrics, Vanderbilt University Medical Center, Nashville, Tennessee, 37232, USA; Imagine Pharma, Devon, Pennsylvania, 19333, USA; Institute of Cellular Therapeutics, Allegheny-Singer Research Institute, Allegheny Health Network, Pittsburgh, Pennsylvania, 15212, USA; Department of Pediatrics, College of Medicine, University of Florida, Gainesville, Florida, 32610, USA; Vanderbilt Program in Developmental Biology, Vanderbilt University, Nashville, Tennessee, 37232, USA; Department of Molecular Physiology and Biophysics, Vanderbilt University, Nashville, Tennessee, 37232, USA; VA Tennessee Valley Healthcare System, Nashville, Tennessee, 37212, USA

## Abstract

Human endocrine cell differentiation and islet morphogenesis play critical roles in determining islet cell mass and function, but the events and timeline of these processes are incompletely defined. To better understand early human islet cell development and maturation, we collected 115 pediatric pancreata and mapped morphological and spatiotemporal changes from birth through the first ten years of life. Using quantitative analyses and a combination of complementary tissue imaging approaches, including confocal microscopy and whole-slide imaging, we developed an integrated model for endocrine cell formation and islet architecture, including endocrine cell type heterogeneity and abundance, endocrine cell proliferation, and islet vascularization and innervation. We also assessed insulin and glucagon secretory profiles in isolated islet preparations from pediatric donors aged 2 months to 10 years and found a temporal difference in the maturation of insulin secretion compared to glucagon secretion. This comprehensive summary of postnatal and pediatric pancreatic islet development provides a framework for future studies and integration of emerging genetic and genomic data related to islet biology and diabetes risk.

## INTRODUCTION

The reduction of functional β cell mass and the failure of the pancreatic islet to adequately regulate blood glucose homeostasis are foundational to all forms of diabetes. Human β cell mass varies among individuals by up to 40-fold during the prenatal period^1–5^ and 3- to 5-fold in adults^6^. The composition of adult human islets also shows considerable heterogeneity among individuals^7^ . Although incompletely defined, insulin secretion by human β cells varies between islets isolated from pediatric and adult pancreas^8^. Collectively, such variations likely affect an individual’s risk and timeline for developing diabetes, yet significant gaps exist in our understanding of the spatiotemporal dynamics of β cell mass, islet cell composition, and function during prenatal and postnatal pancreas development. Furthermore, the β cell-directed autoimmunity of type 1 diabetes often appears in the first five years of life^9,10^ coincident with an active period of pancreas and islet growth and development involving significant tissue reorganization and maturation^1,11,12^.

Defining human prenatal to early postnatal islet cell development has been challenging due to limited access to human tissue at these stages, along with difficulties visualizing islet developmental processes and cellular relationships, both at high resolution and broader overview, in the context of a rapidly growing and reorganizing organ. Histological analyses of human prenatal tissues have shown that the pancreatic epithelium undergoes active growth and tubular morphogenesis around 6-7 weeks gestation (G6-7w)^13,14^. At this point, cells express several key transcription factors such as pancreas and duodenum homeobox 1 (PDX1), sex-determining region Y (SRY) box 9 (SOX9), Nirenberg and Kim homeobox 6.1 (NKX6.1), GATA transcription factor 4 (GATA4), and forkhead box A2 (FOXA2)^3,13,15,16^, which closely resembles the signature of murine multipotent pancreatic progenitor cells (MPCs). Using whole-mount labeling and 3-D rendering approaches on intact pancreatic tissue samples from the first trimester, clusters of insulin (INS)-positive cells were found to be preferentially located in the center of the pancreas, aligned along the longitudinal axis, and essentially absent from the periphery^17^. Ensuing developmental steps that define tip-trunk compartmentalization of epithelial tubes, endocrine cell birth, and islet formation, including the degree to which these processes are synchronized across different pancreatic regions with organ expansion during later stages of prenatal development and after birth, are largely uncharacterized in the human pancreas.

Most postnatal human islet development studies have focused on β cell proliferation and maturation^5,8,18^. Young β cells are quite proliferative during the gestational and perinatal periods, but their proliferative capacity declines sharply in the years after birth, along with their responsivity to cellular mitogens^1,3,5,19,20^. Because postnatal β cell expansion is thought to be primarily achieved through replication, this brief window of proliferative capacity is of great interest to those developing regenerative therapies to replace or restore β cell mass in diabetes. Furthermore, an initial lower β cell mass in this early postnatal period may render an individual more susceptible to diabetes. Also unknown is when glucose-stimulated insulin secretion (GSIS) in humans starts to develop, as GSIS is not present prenatally^19,20^ and pediatric islets have reduced basal insulin secretion compared to adult islets^8^. Likewise, it is unknown when human islets acquire the essential counter-regulatory mechanisms controlling glucagon secretion. Islet vascularization and innervation have been described during prenatal human development^21–25^ but have not been extensively characterized postnatally. Based on studies of mouse pancreas development^26–28^, these processes likely influence endocrine cell maturation in a reciprocating manner.

Extensive remodeling in the cellular architecture and organization of islets after birth and throughout the pancreas has been described^5,29,30^. More recently, cellular polarity, 3D compaction, and the arrangement of mouse islets have been shown to be important in the functional maturation of the islet unit^31^. Additional studies have demonstrated that β cells undergo epigenetic remodeling as they age, substantially affecting their function^32,33^. These studies suggest there are dynamic islet structural and functional changes during postnatal development, but the timing and sequence of these changes remain poorly defined.

Here, we assembled the largest cohort of human pediatric pancreas samples analyzed to date – 115 organs spanning birth through the first decade of life – and systematically applied a combination of complementary tissue imaging approaches, including confocal microscopy and whole-slide imaging to document key morphogenic events of postnatal pancreas and islet development. We generated an immunohistochemistry-based spatiotemporal map of islet formation, quantification of endocrine cell birth, proliferation, and composition, and the acquisition of islet innervation, identifying critical differences between human and rodent islet development. Finally, we characterized pediatric islet function in a perifusion system and showed that islets in the first two years of life have distinct differences in insulin and glucagon secretion dynamics compared to older pediatric and adult islets. Together, the data reported herein provide an unprecedented overview of human islet development that can inform future disease-specific studies of endocrine cell differentiation and function, ultimately leading to the development of new diabetes treatments and paving the way to disease prevention.

## RESULTS

### Collection of large pediatric pancreas cohort through nationwide partnerships with organ procurement organizations

The mortality rates of infants below one year of age and children aged 1-4 years are low in the United States (543.6 and 25.0 deaths per 100,000 population, respectively)^34^, and clinical transplantations are necessarily prioritized from organ donations. Therefore, a very limited number of pediatric pancreata become available for research each year (3-20 for age <1 year; 11-27 for age 1-5 years; 21-33 for age 6-10 years; Organ Procurement and Transplantation Network national data 2014-2023, https://optn.transplant.hrsa.gov/). Building on the infrastructure developed by the Network for Pancreatic Organ Donors with Diabetes (nPOD; RRID:SCR_014641, http://www.jdrfnpod.org) and their outreach to a nationwide network of organ procurement organizations (OPOs) and nonprofit organizations, our team coordinated the collection of over 100 postnatal pediatric organs for research since 2012.

To ensure accurate tracking of morphological landmarks between tissue samples, coronal sections were taken from the head, body, and tail regions (**Figure 1A**). Acknowledging that developmental stages are likely a continuum, donors were stratified to define critical biological processes over time. Approximately corresponding to key postnatal developmental milestones, we used the following categories to stratify our cohort: neonatal, infancy, and childhood.

**Figure 1.**
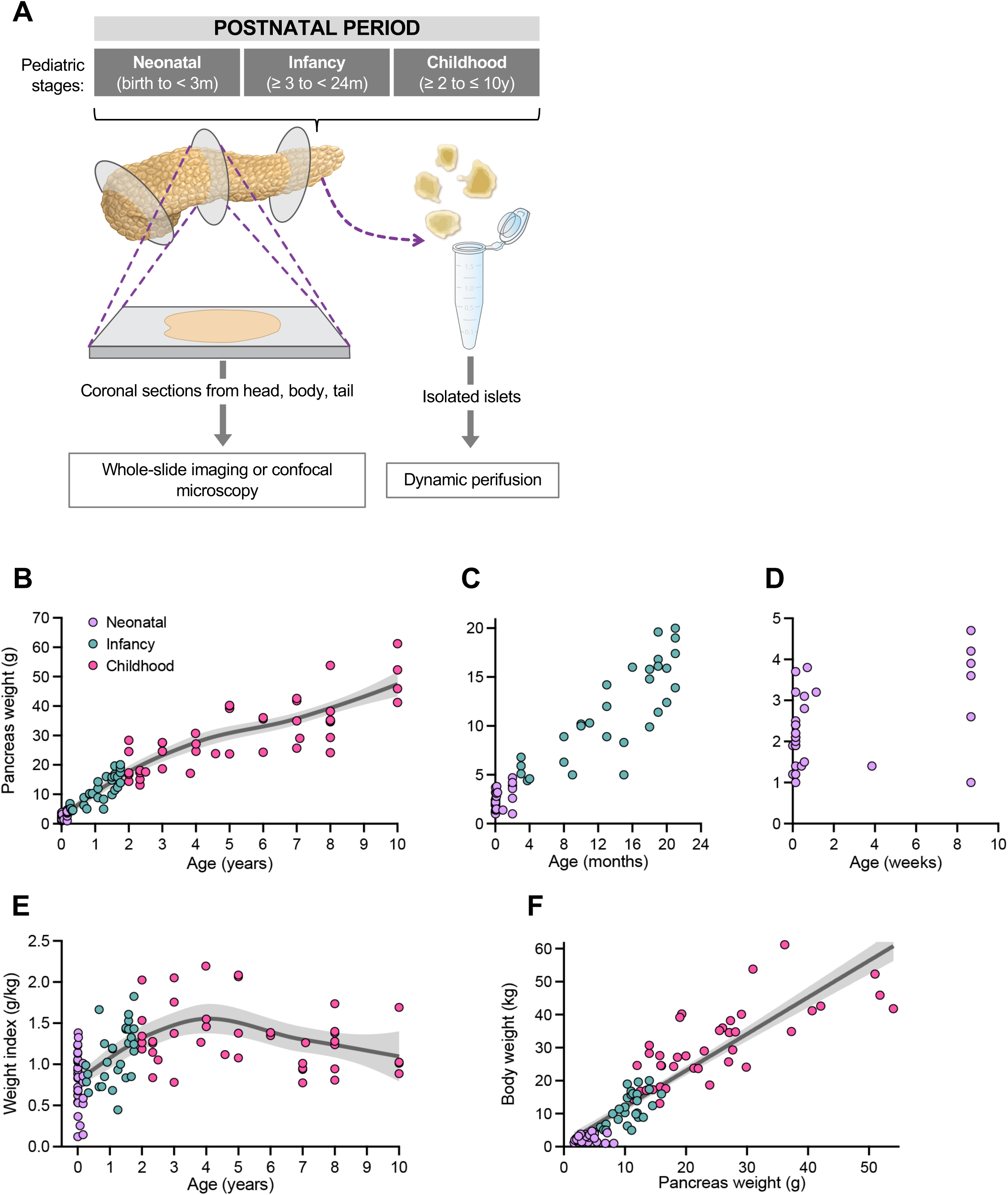
Coordinated analyses of tissue and islets from postnatal pediatric pancreas enabled study of endocrine cell birth, distribution, and function during rapid organ growth across three postnatal stages. (A) Schematic of postnatal pancreas sectioning and analysis. For pancreata from donors aged birth to 10 years, coronal sections were taken from head, body, and tail. Cryosections from all samples were labeled via immunofluorescence using hormone and Ki67 markers, then imaged and analyzed using a cellular recognition algorithm. For a subset of donors, tissue was also digested for islet isolation. (B–D) Pancreatic weight increased with age. Gross pancreatic weights (g) of donor pancreata from neonatal (birth to <3 months, *n*=29), infancy (≥3 months to <24 months, *n*=30), and childhood (≥2 years to ≤10 years, *n*=41) developmental stages were recorded prior to sample processing. C–D are insets of graph in B to illustrate weight range from neonatal and infancy stages. (E) Pancreatic weight index (ratio of pancreatic weight to body weight) increased in a nonlinear fashion with increased age. (F) Pancreatic weight correlated with body weight (Pearson’s product-moment correlation, R=0.899, *p*<0.0001). Panels B and E-F show best fit curve or line ± 95% confidence interval.

Although “neonate” is a term usually reserved for babies four weeks of age or younger^35^, because of the high proportion of anencephalic cases in our cohort (*n*=22, aged 0-5 days), we expanded our definition of neonatal to include donors younger than months of age (totaling *n*=35; see **Table S1**). The infancy stage (*n*=35), defined in this study as ≥3 months to <24 months, corresponds to a period of heightened growth and transitioning nutritional demands^36,37^. The childhood stage (*n*=50) encompassed ages ≥2 years to ≤10 years.

We recognize that the breadth, impact, and information of any imaging data cannot be fully communicated through a limited number of static figures published within journal article constraints. Therefore, images presented herein are deposited in the publicly available Pancreatlas™ platform^38^, which allows the exploration of full-resolution pancreas imaging data in an interactive fashion (https://pancreatlas.org/datasets/531/explore).

As shown in **Figures 1B-D** and **S1**, pancreas weight at birth varied almost 4-fold (0.98 to 3.71g; *n*=19) but showed a rapid, nonlinear increase with increasing postnatal age (F=10.98, *p*=2.74e-6). Because overall body weight varied substantially among individuals of the same postnatal age, pancreas weight was alternatively expressed as a “weight index” (in grams, over total body weight, in kilograms; **Figure 1E**). Interestingly, although weight index showed a nonlinear trend as a function of age (nonlinearity test, F=11.81, *p*=1.21e-6) and means differed among age groups (nonlinearity test, F=22.68, *p*=1.1e-8; **Figure S1D**), standard deviations were not significantly different (Brown-Forsythe test, F=0.06396, *p*=0.9). This similar trajectory of weight index values across neonatal, infancy, and childhood groups suggests that the variability in pancreas size previously noted in adolescents and adults^39,40^ might already be established by birth. Importantly as the cause of death in many neonatal cases was anencephaly (**Table S1**), our studies demonstrated that there was no significant difference in the pancreas weight index when compared with non-anencephalic cases of similar age (**Figure S1E**). Across all postnatal stages, body weight was highly correlated with pancreas weight (Pearson’s product-moment correlation, R=0.899, *p*<0.001; **Figure 1F**).

### Human multipotent progenitors and proto-endocrine cells persisted after birth

During the neonatal period, the pancreatic epithelium maintained a tubular structure. The marker GP2, which identifies multipotent progenitor cells (MPCs)^41^, was prominently expressed at the edges of pancreatic lobes at birth and in the presumptive acinar tissue, but it was absent from endocrine cell clusters (**Figures 2A** and **S2A-B**). This finding is consistent with previous reports showing GP2 expression in pancreatic MPCs, which is downregulated in differentiated endocrine and ductal cells^41,42^. We then investigated the compartmentalization of the pancreatic epithelial tubes and visualized the distribution of two key transcription factors, PTF1A and NKX6.1. These factors are known to function as opposing determinants of acinar versus endocrine lineages in MPCs during mouse pancreas organogenesis^43^. As expected, PTF1A was localized to what has been referred to as the “tip” region of the branched epithelial network^43^. In sharp contrast to earlier findings in the developing mouse pancreas^43,44^, we discovered that NKX6.1 expression extended from the “trunk” to the “tip” region and was co-expressed with PTF1A in GP2+ cells (**Figure 2B-C’**). These observations raise the possibility that endocrine cells in the human pancreas continue to be generated de novo after birth, which is supported by the idea of NKX6.1 functioning as a transcriptional repressor of PTF1A by binding to the PTF1A enhancer^43,45^. After identifying PTF1A+ NKX6.1+ cells in the postnatal pancreas (**Figures 2B-C’**), we also observed proto-endocrine PAX6+ post-mitotic cells that had not yet differentiated regarding hormone specificity, even up to 20 months of age (**Figures 2D-E’** and **S2C-C’**). These findings indicate that the peripheral regions of pancreatic lobes retain residual MPCs and their derived late-born endocrine cells. Together, these data highlight a crucial distinction in human pancreatic endocrine cell development compared to mouse models^44,46,47^: multipotent progenitor cells persist postnatally, suggesting that endocrine cell neogenesis continues after birth.

**Figure 2.**
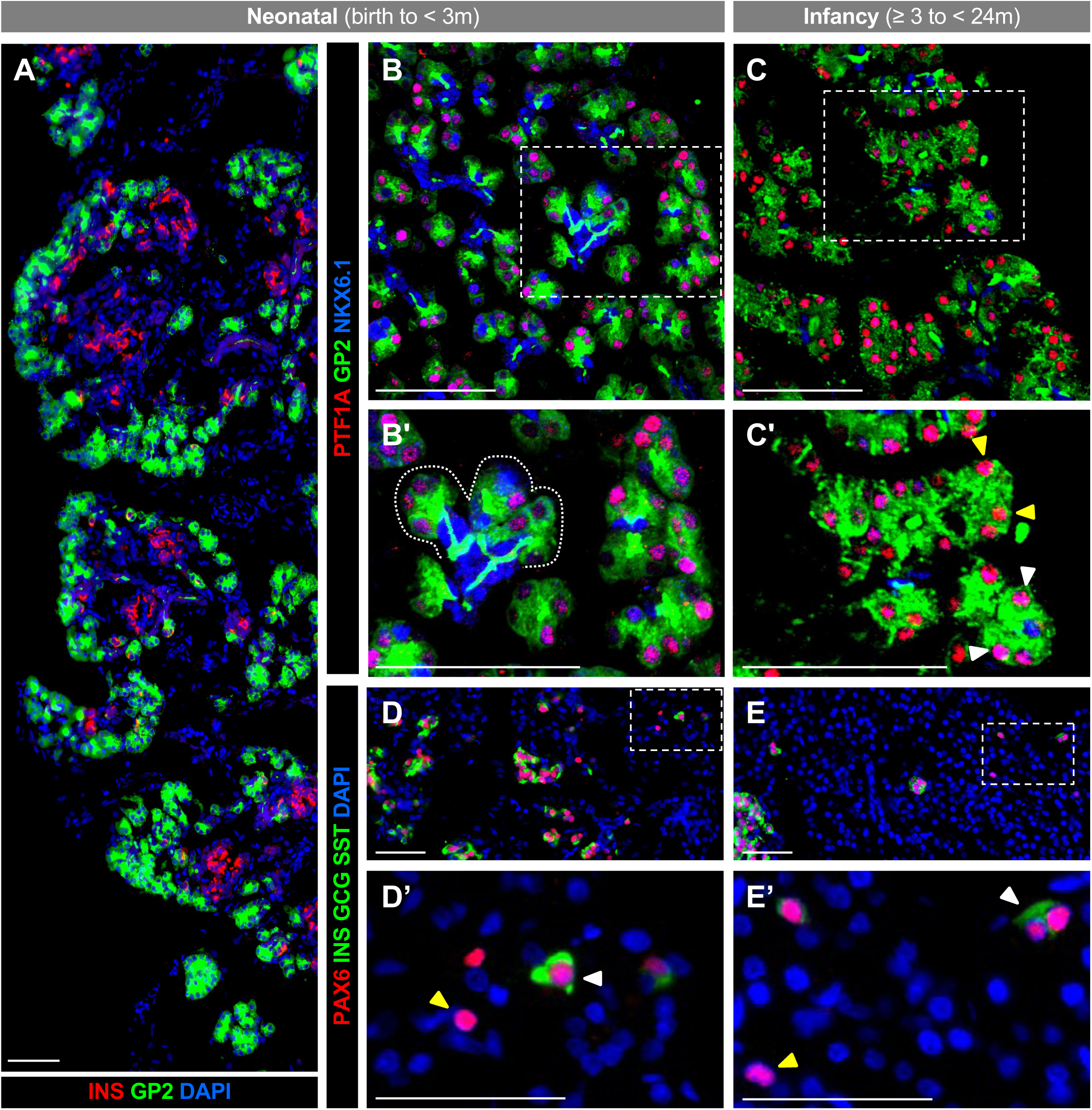
GP2+ multipotent progenitors, as well as proto-endocrine PAX6+ post-mitotic cells, persisted in human pancreas after birth. **(A)** Immunohistochemical staining of pancreatic section from human neonatal donor (1 day) born at 37 weeks of gestation. GP2 expression was highest around periphery of pancreatic lobes. **(B–C’)** Multipotent progenitor cells expressing GP2 (green), PTF1A (red), and NKX6.1 (blue) persisted after birth. Tip regions are outlined by dotted line; areas within boxes marked by dashed line are magnified in **B’–C’**. **(D–E’)** Newly formed endocrine cells appeared around lobe periphery during the neonatal and infancy stages. Areas within boxes marked by dashed line are magnified in **D’–E’**; white arrowheads indicate PAX6+, hormone-positive cells; yellow arrowheads indicate PAX6+, hormone-negative cells. All scale bars are 50 μm.

### Postnatal endocrine cell development was characterized by rapidly declining proliferation and an increased β cell proportion

Small endocrine cell clusters were densely packed across the entire pancreas at birth, both in the periphery and central regions of lobes (**Figures 3A-A’** and **S3**). By infancy, larger clusters of endocrine cells were observed with distinct morphology reminiscent of rodent islets (**Figure 3B-B’**). By childhood, islets were generally indistinguishable from adult islets in structure and composition and more sparsely distributed throughout exocrine tissue (**Figure 3C-C’**), as quantified in **Figure 3D** and **Figure S4A**, showing that the total endocrine cell fraction was high (accounting for 11.63 ± 0.64% of total cells, *n*=24) at birth but rapidly declined thereafter. When considering the endocrine cells alone, proportions of α, β, δ, ε, and γ cells were remarkably consistent among donors of the same age group, with the most striking change occurring between the neonatal and infancy stages (**Figure 3E**). In neonatal pancreata (*n*=24), β cells were the most abundant endocrine cell subtype (42.38 ± 1.75%) followed by δ cells (34.04 ± 1.50%), then α cells (23.58 ± 1.31%). In the infancy stage (*n*=21), the proportion of β cells was increased (55.30 ± 1.81%), while the α cell proportion remained unchanged (21.98 ± 1.38%) and the δ cell proportion decreased (22.72 ± 0.87%). These β, α, and δ proportions were maintained in childhood at 57.89 ± 1.00, 23.25 ± 0.81, and 19.05 ± 0.62%, respectively. The numbers of ghrelin-producing ε cells and pancreatic polypeptide-producing γ cells remained consistently low (<6% each, *n*=4 per stage) throughout neonatal, infancy, and childhood stages.

**Figure 3.**
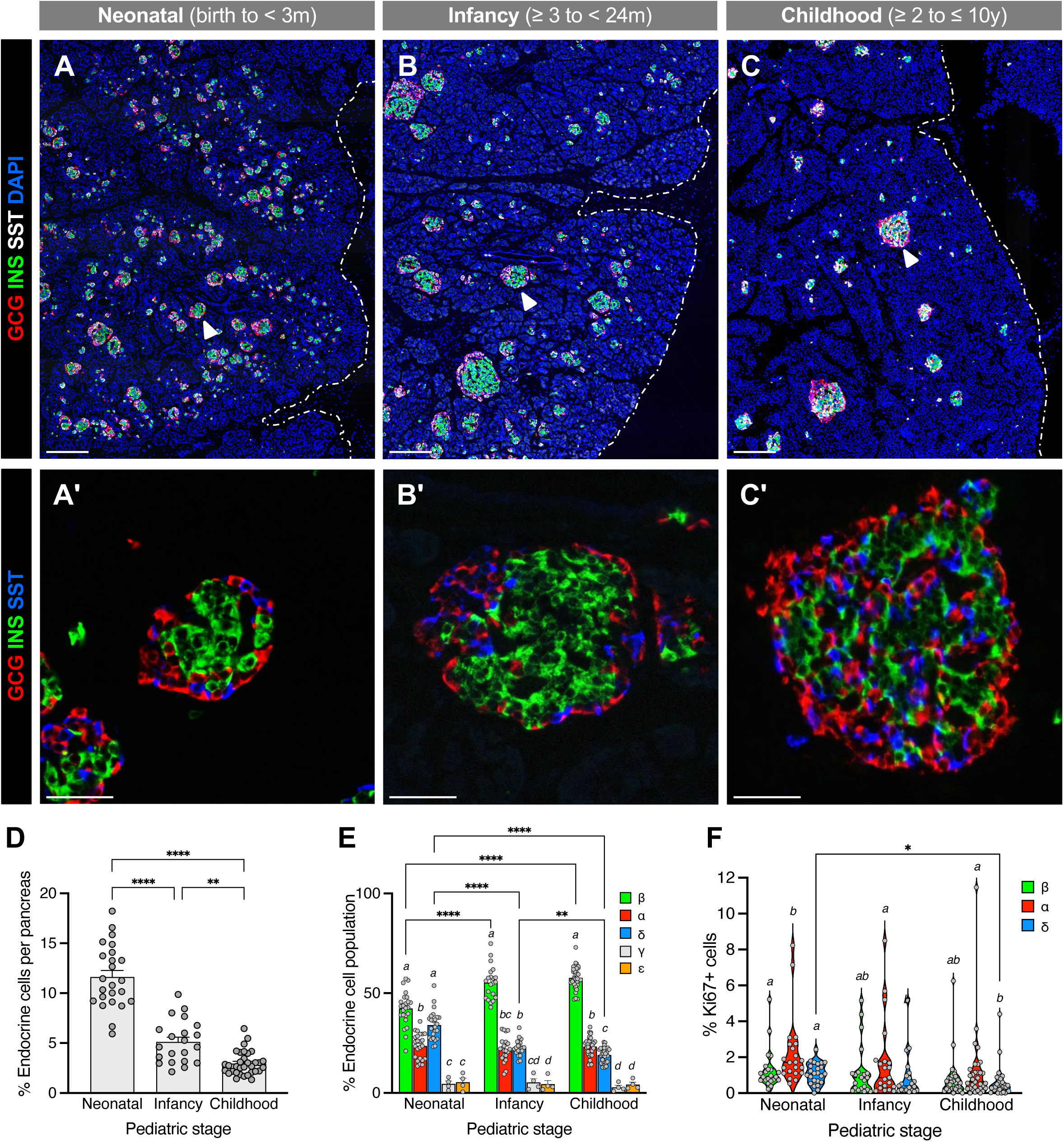
Endocrine cell density and proliferation rates rapidly declined during postnatal development as the β cell population expanded in proportion. **(A–C)** Pancreatic sections showing islet distribution during neonatal (**A**), infancy (**B**), and childhood (**C**) stages. All scale bars are 200 μm. **(A’–C’)** Representative islets (white arrowheads) from each stage are shown at higher magnification. All scale bars are 50 μm. **(D–F)** Quantification (mean + SEM) of endocrine cell density (**D**), islet composition (**E**), and endocrine cell proliferation (**F**); neonatal, *n*=26; infancy, *n*=22; childhood, *n*=33. Data are plotted as mean + SEM (**D**–**E**) or median ± quartiles (**F**), with each circle representing one individual. Means were analyzed by one-way ANOVA (**D**) or mixed-effects analysis (**E**–**F**) followed by Tukey’s multiple comparisons tests. Brackets with asterisks denote differences between age groups; *, *p*<0.05; **, *p*<0.01; ***, *p*<0.001; ****, *p*<0.0001. Italicized letters denote comparison among cell types within each age group; bars with like letters do not statistically differ.

To account for the variable growth rates among individuals of similar ages, pancreatic composition was further analyzed as a function of total pancreas weight. Densities of α, β, and δ cells showed significant nonlinearity relative to increasing pancreas weight (**Figure S4B**; F=17.65 α, 11.12 β, 22.90 δ; all *p*<0.001), consistent with the trajectory of exocrine tissue expansion. Strikingly, β and δ cell composition showed nonlinear changes relative to increasing pancreas weight (F=6.67 β, 8.06 δ; both *p*<0.001), while the proportion of α cells remained consistent (approximately 22%; F=1.03, *p*=0.3861; **Figure S4C**) with no linear relationship between α cell numbers and pancreas weight (*p*=0.876). These data indicate that postnatal α cell composition is surprisingly stable while β and δ cell populations exhibit differential populational growth trajectories from birth to childhood.

Postnatal β, α, and δ cell proliferation rates were quite low (less than 2.5%), even during the neonatal stage when endocrine cell density was at its peak (**Figures 3F** and **S4A**). Notably, α cell proliferation (2.25 ± 0.41%) was significantly greater than δ cell proliferation (1.09 ± 0.12%; adjusted *p*=0.0053) and β cell proliferation (1.31 ± 0.24%; adjusted *p*=0.0055) at this stage. All three subpopulations exhibited comparable proliferation rates during infancy (1.22 ± 0.32 β, 1.75 ± 0.48 α, 0.87 ± 0.26% δ) and childhood (0.76 ± 0.19 β, 1.20 ± 0.35 α, 0.58 ± 0.14% δ).

Comparing between neonatal and childhood stages, only δ cell proliferation was significantly decreased (adjusted *p*=0.0215). While previous studies have linked on-ventilator time to increased endocrine proliferation rates^48^, we did not observe such a correlation with donors in this study for whom the data was available (**Figure S4D-E**). Moreover, since the cause of death for many of our neonatal donors was anencephaly, we investigated whether this condition influenced neonatal pancreas or islet composition. When considering anencephalic cases (*n*=22, aged 0-5 days) compared to non-anencephalic neonatal donors (*n*=13, aged 1 day to <3 months), we observed no difference in endocrine density, endocrine cell composition, or endocrine cell proliferation (**Figure S4F-H**). These data suggest that in utero pancreas development occurs normally despite neural tube abnormalities in these cases^49,50^. Overall, our data point to highly variable postnatal endocrine proliferation rates among individuals.

### Islet architecture shifted from a core-mantle structure to more heterotypic cell-cell positioning by two years of age

In addition to proportional changes in islet cell composition during pancreas growth, spatial alterations were also apparent. Just after birth, β cells were found both in the islet periphery (defined as within 20 μm from outermost cells; **Figure 4A**) and deeper in the islet “core” (beyond 20 μm), with slightly more β cells in the core (55% vs. 45%, **Figure 4B**). By childhood, β cell distribution had shifted to 62% of β cells localized to the islet periphery and 38% in the core. In contrast, both α and δ cells were overwhelmingly located in the islet periphery just after birth (91% and 88% of all α and δ cells, respectively) with these percentages decreasing somewhat by childhood (72% α and 59% δ; **Figure 4C-D**). As a result, α, β, and δ cell types were considerably more intermingled by childhood (**Figure 4C’**) compared to the prominent β cell “core” seen in the first two years after birth (**Figure 4A’-5B’**).

**Figure 4.**
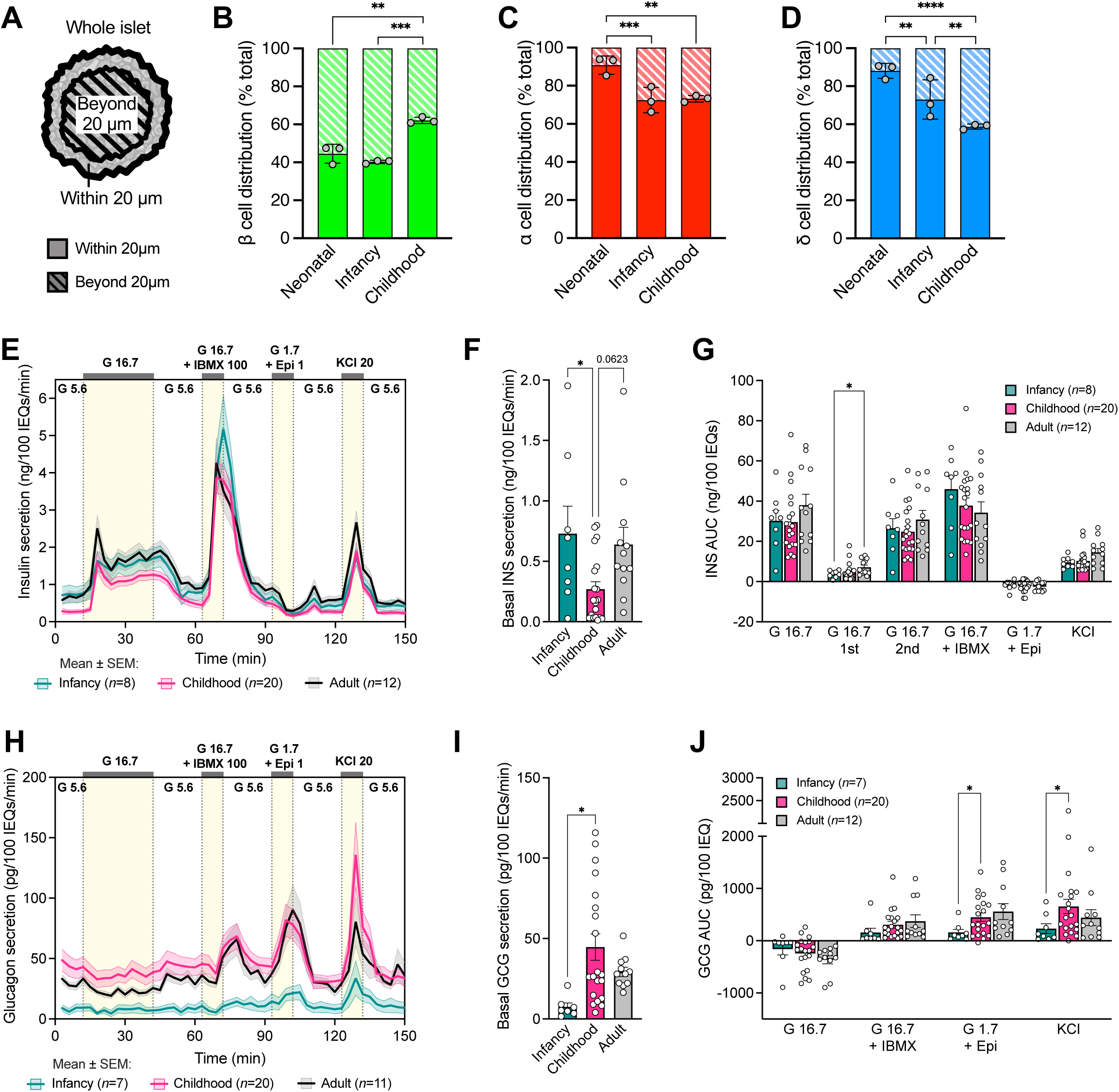
Human islets gradually shifted away from murine-like arrangement of β cell-rich “core” concomitant with maturation of functional glucagon secretory response. **(A)** Schematic depicting quantification of β, α, and δ cell distribution within the first 20 μm of the islet border (periphery) and beyond 20 μm depth (interior). **(B–D)** Distribution of β, α, and δ cells between islet periphery vs. interior across three postnatal stages (*n*=3 donors per stage); symbols represent the mean of 30 islets per donor. Standard error shown per stage. **(E**–**J)** Dynamic insulin (**E**–**G**) and glucagon (**H**–**J**) secretory response of islets from infancy (*n*=8), childhood (*n*=20), and adult (*n*=12) independent donors measured by perifusion, normalized to islet volume. Differences in secretion trajectories normalized to islet volume were analyzed by linear mixed-effect models for insulin (**E**, global *p*=1.094e-37; pairwise comparisons infancy vs. childhood, *p*=2.860e-27; infancy vs. adult, *p*=6.085e-20; childhood vs. adult, *p*= 6.445e-32) and glucagon (**H**, global *p*=3.059e-23; p=2.054e-15; pairwise comparisons infancy vs. adult, *p*=2.441e-14; childhood vs. adult, *p*= 8.798e-17). Panels **G** and **J** show secretory response as calculated by area under the curve (AUC). Treatments (and concentrations) are shown along the top of the graph: Epi, epinephrine (μM); G, glucose (mM); IBMX, isobutylmethylxanthine (μM); KCl, potassium chloride (mM). Means were analyzed by one-way (**F, I**) or two-way (**B–D**, **G**, **J**) ANOVA followed by Tukey’s multiple comparisons tests. Brackets with asterisks denote differences between age groups; *, *p*< 0.05; **, *p*<0.01; ***, *p*< 0.001; ****, *p*< 0.0001.

**Figure 5.**
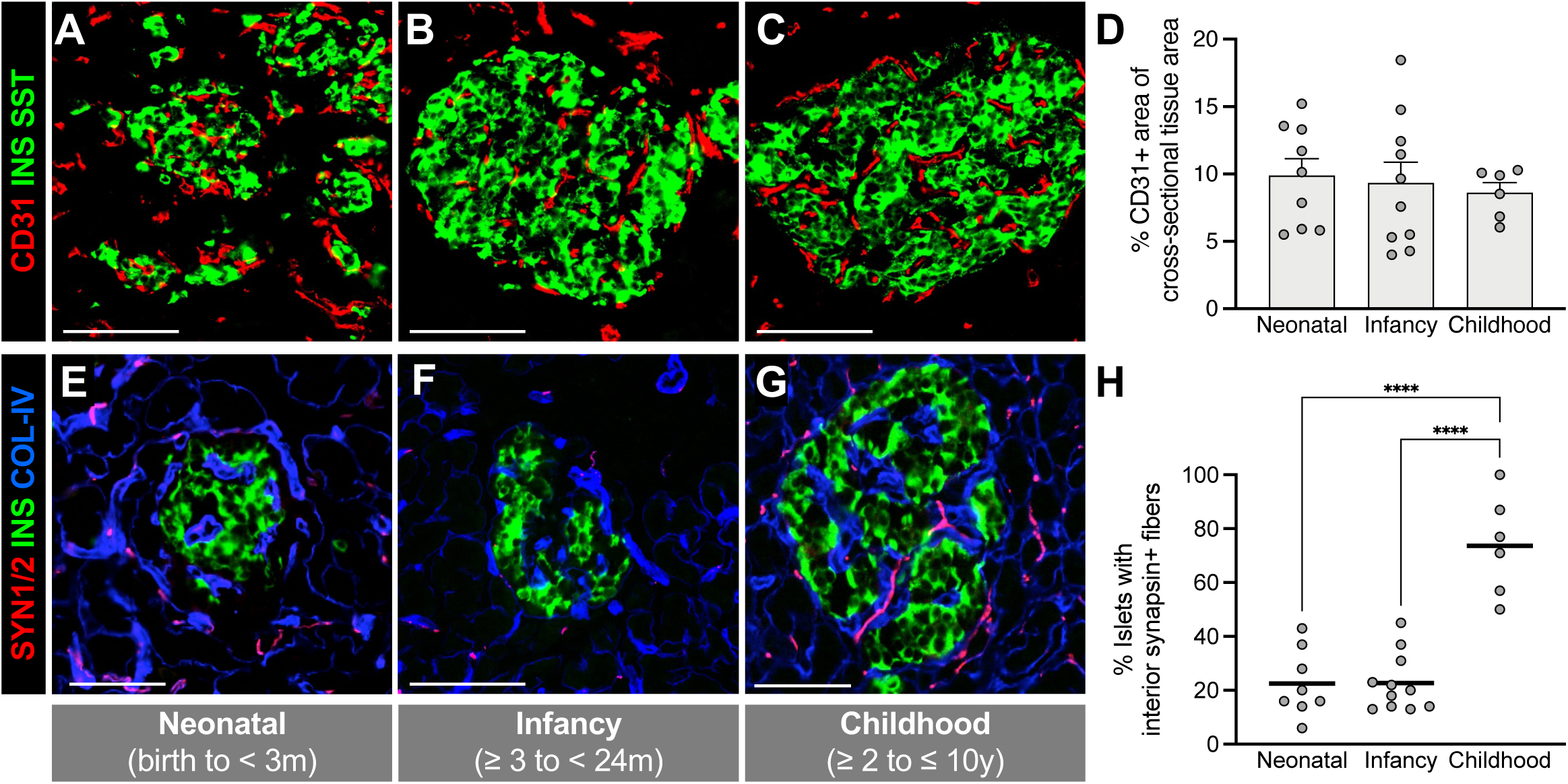
Islet innervation was delayed compared to islet vascularization. Asynchronous vascularization (**A-C**; defined as percentage CD31+ area of total cross-sectional tissue area in **D**) and innervation (**E-G**; defined as percentage of islets with interior synapsin-1/2+ fibers in **H**) were quantified across neonatal (*n*=8-9), infancy (*n*=10-11), and childhood (*n*=6) stages. Whereas vascular density was comparable from neonatal to childhood, islets were not innervated at birth, but axons gradually extended to islets postnatally. All scale bars are 100 µm. Means were analyzed by one-way ANOVA followed by Tukey’s multiple comparisons tests. Brackets with asterisks denote differences between age groups; *, *p*< 0.05; **, *p*<0.01; ***, *p*< 0.001; ****, *p*< 0.0001.

### Maturation of glucagon secretion profile occurred later than insulin secretion

Islet isolation from pediatric pancreas donors expanded on previously published work^51^ and incorporated extensive prior experience in isolating islets from organ donors with type 1 diabetes^52^, type 2 diabetes^53^ and cystic fibrosis-related diabetes^54^. This allowed us to optimize islet isolation and tissue processing of young human pancreata with the aim to consistently obtain fractions of free, non-mantled islets for islet functional analysis. We performed dynamic perifusion experiments on islets isolated from donors in infancy (*n*=8; ages 2-19 months for functional studies) and childhood (*n*=20; ages 2-10 years) stages and compared the responses to islets from adult donors (*n*=14; ages 24-59 years) isolated at the same islet isolation facility (**Figures 4E-J** and **S5A-K**). Basal insulin secretion was significantly lower in childhood compared to infant islets when normalized to islet equivalents (IEQ) (**Figure 4E-F**) but did not reach statistical significance when secretion was normalized to insulin content (**Figure S5A-C**). An earlier study found that basal insulin secretory response normalized to islet volume (IEQ) appeared to be lower in postnatal pediatric islets (<10 years of age) compared to adult, but donor islets were not stratified into infancy and childhood categories due to a small sample size^8^. To comprehensively analyze insulin secretory response in the presence of different secretagogues, we employed a linear mixed-effects model^55^ that accounts for underlying temporal correlations and demonstrated global statistically significant differences in insulin secretory trajectories across the three age groups of infancy, childhood, and adulthood (**Figures 4E** and **S5B**). Subsequent binary comparisons between each pair of age groups also demonstrated statistical significance between trajectories when insulin secretion was normalized to islet volume (**Figure 4E**) or insulin content (**Figure S5B**). In contrast, analysis of insulin secretory response by area under the curve (AUC) normalized to IEQ – representing total insulin secreted in response to a given stimulus – revealed no significant differences between age groups under glucose-stimulated, cAMP-potentiated, or KCl depolarization-mediated conditions (**Figure 4G**). However, first-phase GSIS was lower in infancy-stage islets compared to adult (**Figure 4G**).

Like insulin, glucagon secretory trajectories analyzed by a linear mixed-effects model revealed statistically significant differences among infancy, childhood, and adult stages (**Figures 4H** and **S5F**). Basal glucagon secretion normalized to IEQ was significantly greater in childhood compared to infancy (**Figure 4I**) and low glucose/epinephrine-stimulated and KCl-mediated glucagon measured by AUC were significantly greater in childhood compared to infancy (**Figure 4J**). However, these differences were not discernible when normalizing glucagon secretion to glucagon content (**Figure S5E-H**). No significant differences were observed in cAMP-potentiated glucagon secretion or high glucose-mediated glucagon suppression (**Figures 4J** and **S5H**).

Due to changing morphological features across pediatric stages, we also considered whether any functional parameters in our study were associated with age as a continuous variable. There was no statistically significant correlation between islet hormone content and age; however, basal insulin and glucagon secretion normalized to IEQ were negatively and positively associated with age, respectively (Spearman correlation; *r*=–0.455, approximate *p*=0.0148, insulin; *r*=0.617, approximate *p*=0.0006, glucagon) (**Figure S5K**). Overall, data point to qualitative and quantitative changes in hormone secretory profiles during postnatal development compared to the adult stage.

### Formation of intra-islet capillary network preceded islet innervation

Vascular endothelial cells have previously been observed adjacent to endocrine cells during prenatal development^23,24,29,56,57^. While neuronal fibers associate with larger vascular structures, they are reportedly absent in prenatal endocrine cell clusters^5,22,25,58^. This prenatal vascular pattern persisted during the early postnatal period, and the overall cross-sectional area occupied by CD31+ vascular structures was similar across neonatal, infancy, and childhood stages (**Figures 5A-D** and **S6A**). At birth and in the first two years of life, the pattern of nerve fibers in islets was variable among donors (10-40% islets with interior synapsin+ fibers) compared to childhood, where most islets (>60%) had interior nerve fibers (**Figures 5E-H** and **S6B**). Thus, human islet innervation occurs later than islet vascularization, similar to what is seen in mice^28^, although adult human islets are comparatively less vascularized and innervated, while human acinar tissue has higher nerve fiber density^59^.

### Model of dynamic changes throughout prenatal and postnatal islet development

Data in this study support a model of tightly regulated islet formation and endocrine cell maturation spanning the postnatal period of human pancreatic development (**Figure 6**). Connecting this data to prior studies in the prenatal pancreas suggests that by early in the second trimester (G13-18w), endocrine cells are forming within the epithelium at the intra-tip and tip-trunk interface. These small clusters of endocrine cells are scattered throughout the developing pancreas, closely associated with blood vessels but away from nerve fibers, which are more abundant in the exocrine tissue^3,13–17^. Due to relatively high endocrine cell proliferation in the neonatal stage, endocrine cells are abundant in the first months after birth, and some α, β, and δ cell proliferation is sustained. However, the rapidly expanding exocrine tissue results in a precipitous decline in overall endocrine cell density despite ongoing endocrine cell expansion. Furthermore, the neonatal and early infancy stage islets exhibit a core-mantle architecture reminiscent of rodent islets. Islets become innervated with nerve fibers months to two years postnatally, considerably later than they become vascularized. By childhood, endocrine cells have undergone extensive rearrangement (possibly because of other islet cell types such as fibroblasts, pericytes, and macrophages and due to ingrowth of nerve fibers), and islet architecture is very similar to that of adult human islets with intermingled α, β, and δ cells^60–62^. At this point, endocrine cell proliferation rates are quite low with β cells comprising the greatest endocrine cell proportion.

**Figure 6.**
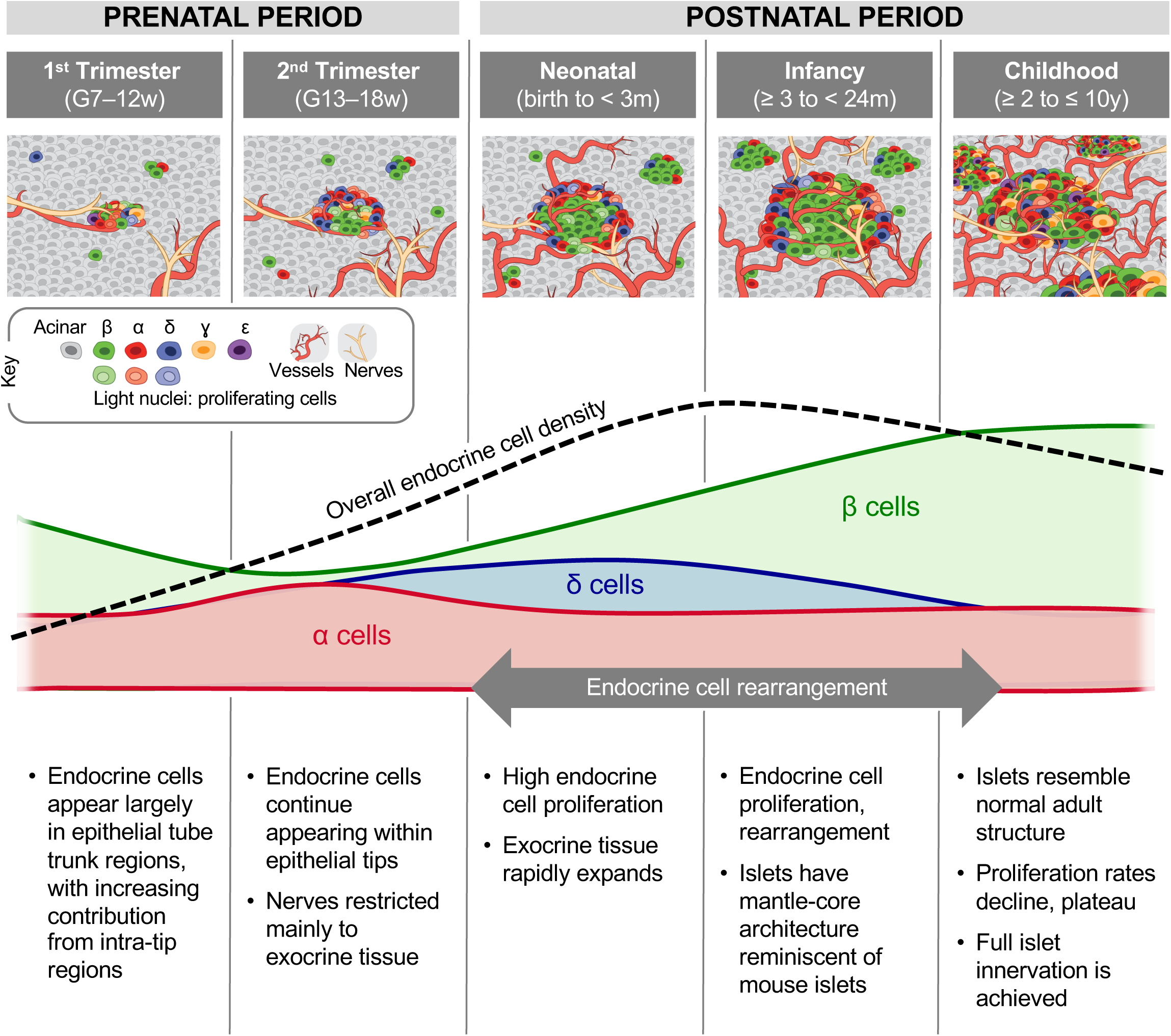
Model describing prenatal and postnatal islet formation. At the onset of 2nd trimester during the prenatal period, most endocrine cells originated from intra-tip and tip-trunk interface. Small clusters of endocrine cells, including PP-rich regions, were found in the core of pancreatic lobes, and nerves were more abundant in the exocrine tissue. The information regarding the development of the prenatal pancreas was obtained from earlier studies^1,3,10,13,16,18^. The neonatal stage was marked by high endocrine cell proliferation and rapid expansion of exocrine tissue, with elevated proliferation sustained during infancy. At this time islets exhibited a core-mantle structure reminiscent of mouse islets. During infancy to childhood stages, endocrine cells underwent extensive rearrangement to resemble adult human islets, and endocrine cell proliferation rates rapidly declined. By this time, islet innervation was achieved.

## DISCUSSION

Human pancreatic endocrine cell differentiation and islet maturation form the basis for islet cell mass and function, which are crucial for the maintenance of normal glucose homeostasis. These dynamic processes begin in utero and continue into early adult life but are incompletely defined, despite likely contributing toward the lifetime risk for all forms of diabetes. Our current understanding is primarily extrapolated from rodent data, which does not sufficiently mimic human development, or from very small studies limited by challenges in acquiring human tissues in this age range. To address these knowledge gaps, this work mapped morphological and spatiotemporal changes in the human pancreas from an unprecedented cohort of 115 organ donors without diabetes, spanning from birth through the first ten years of life. Using quantitative analysis of human islet cells, we identified key steps, insights, and timelines for islet cell development and outlined a comprehensive summary bridging prior work on the prenatal period with our postnatal observations (**Figure 6**). We also assessed dynamic islet function using isolated islet preparations from pediatric organs and identified a different timeline for the maturation of insulin and glucagon secretory profiles.

### Variations in human pancreas size, islet structure, and islet cell composition appeared early in life

In our study of human pancreata from over 100 donors under the age of ten, identified through a national network and processed promptly after death using standardized methods, we discovered that variation in pancreas size among nondiabetic individuals begins as early as neonates and continues during the first months of life. Notably, pancreas weight can differ nearly fourfold at birth. As expected, pancreas weight increased with age and correlated with body weight. These findings strongly suggest that pancreas weight, a potential biomarker for increased risk of type 1 diabetes, is primarily determined prenatally by the intrauterine environment and genetics. A role for genetics in determination of pancreas size was supported by a recent genome-wide association study identifying loci that correlate with pancreas size^63^.

We also found variable endocrine cell abundance among donors at all stages of postnatal pancreas development, suggesting that endocrine cell lineage allocation varies among individuals and is defined early in prenatal development. A similar variation in islet cell composition has been noted in adult human islets^7^. Despite this variability, relative proportions of α, β, and δ cells seemed to reach an equilibrium quickly after birth, with β cells predominating all other endocrine cell populations by three months of age. These results underscore previous studies showing that relative δ cell abundance decreases concomitantly with increased β cell abundance^5,64,65^. Furthermore, the arrangement of endocrine cells within an islet was also dynamic, with islets in the first two years of life having a core of β cells and a mantle of α and δ cells, reminiscent of rodent islets, and then transitioning to the intermingled cell arrangement seen in adult human islets. Additional studies are needed to identify the mechanisms responsible for these changes in islet cell neighborhoods.

### Human islet cell proliferation varied with age, islet cell type, and among individuals

Postnatal endocrine cell development is characterized by rapidly declining proliferation rates and an increase in β cell proportion^1,3,5,19,20^. We extend these prior observations, finding similar β and δ cell proliferation rates across neonatal, infancy, and childhood stages; however, we noted greater α cell proliferation during the neonatal stage with considerable variation among individual donors. These findings suggest that the variation in adult β cell mass has its roots in this early phase of life and that mechanisms other than cellular replication contribute to β cell mass postnatally. Postnatal β cell proliferation rates were low, suggesting that the cellular senescence observed in adult β cells is acquired during the first decade of life. The drivers of this process are largely unknown but may involve ligands such as endocannabinoids, noncanonical Notch signaling, and semaphorin–neuropilin pathways^66–68^.

### Postnatal processes in human endocrine cell neogenesis and islet innervation

Identification of presumed MPCs, cells co-expressing PTF1A and NKX6.1 and localized to epithelial tips, in human postnatal pancreas suggests that endocrine cells are generated by both neogenesis and proliferation but likely at different frequencies across the α, β, and δ cell lineages. MPCs have been of great interest since the initial observation that early progenitor cell proliferation preceded β cell differentiation in grafted human embryonic pancreas^11^, followed by the discovery that single islets contained β cells derived from multiple progenitors^69^. These discoveries informed the development of stem cell differentiation protocols that have enabled the analysis of chromatin, RNA, and proteomic signatures as well as the modulation of external factors that may affect endocrine cell allocation^70–73^. Since then, MPC-associated markers have been identified^41,74,75^ and studied to reconstruct transcriptional trajectory^56,73,76–78^. While spatial resolution and donor number was limited, single cell RNA-sequencing combined with spatial transcriptomics in the prenatal human pancreas identified several novel genes in these early endocrine cell clusters and surrounding microenvironment that may play a role in directing endocrine cell differentiation^78^. Our results suggest that these processes are highly relevant even after birth and thus, may be amenable to modulation or intervention. Additional studies into these MPC-associated markers and genes, along with the identification of additional regulators of prenatal and postnatal endocrine cell differentiation, are needed to clarify the mechanisms that regulate this process. Similar to observations in mouse studies^79^, the vascularization and innervation of islets in humans are not synchronized during development. While human islets are vascularized at birth, innervation occurs later than vascularization. Interestingly, adult human islets are generally less vascularized and less innervated compared to mouse islets^80^. In contrast, human acinar tissue exhibits a higher density of nerve fibers, indicating species-specific differences in the neurovascular patterning of the exocrine pancreas compartment and islets. Additionally, because human islets have fewer autonomous nerve terminals, paracrine signaling may play a more significant role in cell-cell communication in human islets compared to mouse islets.

### Morphological and functional islet cell maturation were asynchronous

To complement immunohistochemistry-based imaging of developmental processes across pancreatic tissue, we examined the function of isolated islets during infancy and childhood. Insulin secretory responses were more similar among infancy and childhood compared to adult islets, though with subtle shifts in magnitude and baseline secretory response. The appearance of comparable high glucose-stimulated glucagon inhibition during infancy and childhood, in contrast to epinephrine-stimulated and KCl-mediated glucagon secretory responses being significantly reduced during infancy, suggests that counterregulatory responses may be acquired in α cells before complete maturation of glucagon secretory machinery. These data, along with our observation of later innervation of islets, suggest that it takes several years after birth for α cells to achieve functional maturation for regulated glucagon secretion. In contrast, β cells are responsive to diverse stimuli much earlier. The lack of significant differences in insulin and glucagon content between age groups indicates that functional maturation is likely mediated by factors different from those regulating hormone production. Future studies are warranted to identify the mechanisms contributing to functional immaturity and its progression in young human islets.

### Limitations

This report presented cross-sectional data; therefore, the conclusions should be interpreted overall and not as a conclusive longitudinal trajectory for any given individual. The use of in vitro cell differentiation systems or other perturbation methods will be required to experimentally test the observations made. Other studies of human β cell development have similarly used human samples but mostly focused on the prenatal period^3,13,16,19,75,81–84^. An advantage of the current report is that we used a harmonized experimental approach to quantify events in postnatal pediatric samples across a large cohort, supporting the generalizability of the proposed model and timeline (**Figure 6**). However, pre-collection events or exposures (e.g., cold ischemia time, nutritional status, in utero environment) could influence these observations, especially in very young donors. Although approximately two-thirds of the neonatal donors (*n*=22 of *n*=35 total; 63%) had anencephaly, we did not observe any significant differences in the parameters studied.

### Summary, future opportunities, and implications for human diabetes

This work connected postnatal endocrine cell development with key morphogenic events previously described during prenatal development, including endocrine cell birth, islet assembly, and the formation of islet neurovascular connections. Utilizing over 15 different markers, we examined islet architecture, endocrine cell type heterogeneity/abundance, endocrine cell proliferation, and islet innervation by immunohistochemistry. These robust spatiotemporal datasets will be a useful reference for β cell regeneration efforts and embryonic stem cell studies by establishing the timeline for human islet architectural and functional maturation.

Furthermore, quantifying islet developmental changes after birth provides critical information to further study the causes and consequences of autoimmunity associated with type 1 diabetes, which often begins in infancy^9,10^. Indeed, it is tantalizing to speculate that the dynamic changes in human pancreatic islets in early life might initiate or accelerate this autoimmunity, representing a “susceptibility window” of some sort. In addition, defining the prenatal and postnatal factors that determine β cell mass in adolescence and adulthood may identify targets for interventions to amplify or enhance β cell mass across all forms of diabetes.

These findings also provide a framework for human pancreatic endocrine cell and islet development that will allow the integration of emerging genetic and genomic data related to islet biology and diabetes risk. Studies mapping the 3D relationships among endocrine cells and epithelium, along with vascular, lymphatic, and neuronal structures, will be crucial to building a comprehensive molecular and morphologic catalog of human pancreas development. Together, these results serve as a foundation for forthcoming studies using emerging technologies such as multiplex imaging, spatial transcriptomics, proteomics, lipidomics, and genomics to characterize and understand the phenotypic signatures, signaling mechanisms, and functional maturation of islet endocrine cells in the first decade of life.

## ACKNOWLEDGEMENTS

We thank the organ donors and their families for their invaluable donations. In addition, these studies were only made possible through the efforts of numerous Organ Procurement Organizations, the International Institute for Advancement of Medicine (IIAM), and the National Disease Research Exchange (NDRI). We also recognize the essential efforts provided by a series of individuals whose dedication to obtaining quality tissues is especially noteworthy, including Alberto Pugliese, Martha Campbell-Thompson, Carmella-Evans Molina, Maigan Brusko, Clive Wasserfall, Helmut Hiller, Robert O’Flynn, and John Kaddis. We thank Joana Almaça, Alejandro Caicedo, Seung Kim, Yan Hang, Robert Whitener, Sarah Richardson, and Noel Morgan for valuable scientific discussion and ongoing collaborative efforts studying pediatric pancreas development. We recognize Samuel Johnson for his dedicated development and management of Pancreatlas^TM^.

This work was supported by The Leona M. and Harry B. Helmsley Charitable Trust [Human Atlas of the Neonatal Developmental and Early Life-Pancreas (HANDEL-P)], G-2004-03813. These efforts were also supported by Human Islet Research Network (RRID:SCR_014393), the Human Pancreas Analysis Program (RRID:SCR_016202), DK106755, DK123716, DK123743, DK120456, DK104211, DK108120, DK104218, DK112232, DK112217, DK117147, DK129469, DK135017, OD0426640, EY032442, DK020593 (Vanderbilt Diabetes Research and Training Center), Breakthrough T1D (formerly JDRF), and the Department of Veterans Affairs (BX000666). This research was performed with additional support from the Network for Pancreatic Organ donors with Diabetes (nPOD; RRID:SCR_014641), a collaborative type 1 diabetes research project supported by Breakthrough T1D (formerly JDRF) (nPOD: 5-SRA-2018-557-Q-R) and The Leona M. & Harry B. Helmsley Charitable Trust (G-2018PG-T1D053, G-2108-04793), and by the Breakthrough T1D (formerly JDRF) Center of Excellence in New England. The content and views expressed are the responsibility of the authors and do not necessarily reflect the official view of nPOD. Organ Procurement Organizations (OPO) partnering with nPOD to provide research resources are listed at https://npod.org/for-partners/npod-partners/. Whole-slide imaging was performed in the Islet and Pancreas Analysis Core of the Vanderbilt DRTC (DK020593). Confocal microscopy was performed in part through the use of the Vanderbilt Cell Imaging Shared Resource (supported by NIH grants CA68485, DK20593, DK58404, DK59637 and EY08126).

## AUTHOR CONTRIBUTIONS

Conceived and designed the experiments: conceptualization by D.C.S., N.H., F.C.P., M.A.A., C.V.E.W., A.C.P., and M.B.; data curation by D.C.S, C.V.R., A.L.H., JP.C., and M.B.; funding acquisition by M.A.A., C.V.E.W, A.C.P, and M.B. Performed the experiments, including investigation and methodology: D.C.S., N.H., F.C.P., C.V.R., A.L.H., N.I., S.M., B.A.S., C.D., J.D., R.H., R.A., H.D., G.P., A.M., I.K., M.L.B., M.Y., D.L.G., L.D.S., R.B., M.A.A., C.V.E.W., A.C.P., and M.B. Analyzed the data: D.C.S., N.H., F.C.P., C.V.R., A.L.H., N.I., S.M., B.A.S., C.D., J.D., R.H., R.A., H.D., G.P., A.M., H.K., K.I.A. and M.B. Contributed materials and/or analysis tools: resources from D.C.S, JP.C., M.A.A., A.P.C., and M.B. Manuscript writing: visualization by D.C.S. and M.B.; original draft by D.C.S., K.I.A., M.A.A., C.V.E.W, A.C.P., and M.B.; review and editing by all authors.

## SUPPLEMENTARY FIGURE LEGENDS

**Figure S1.**
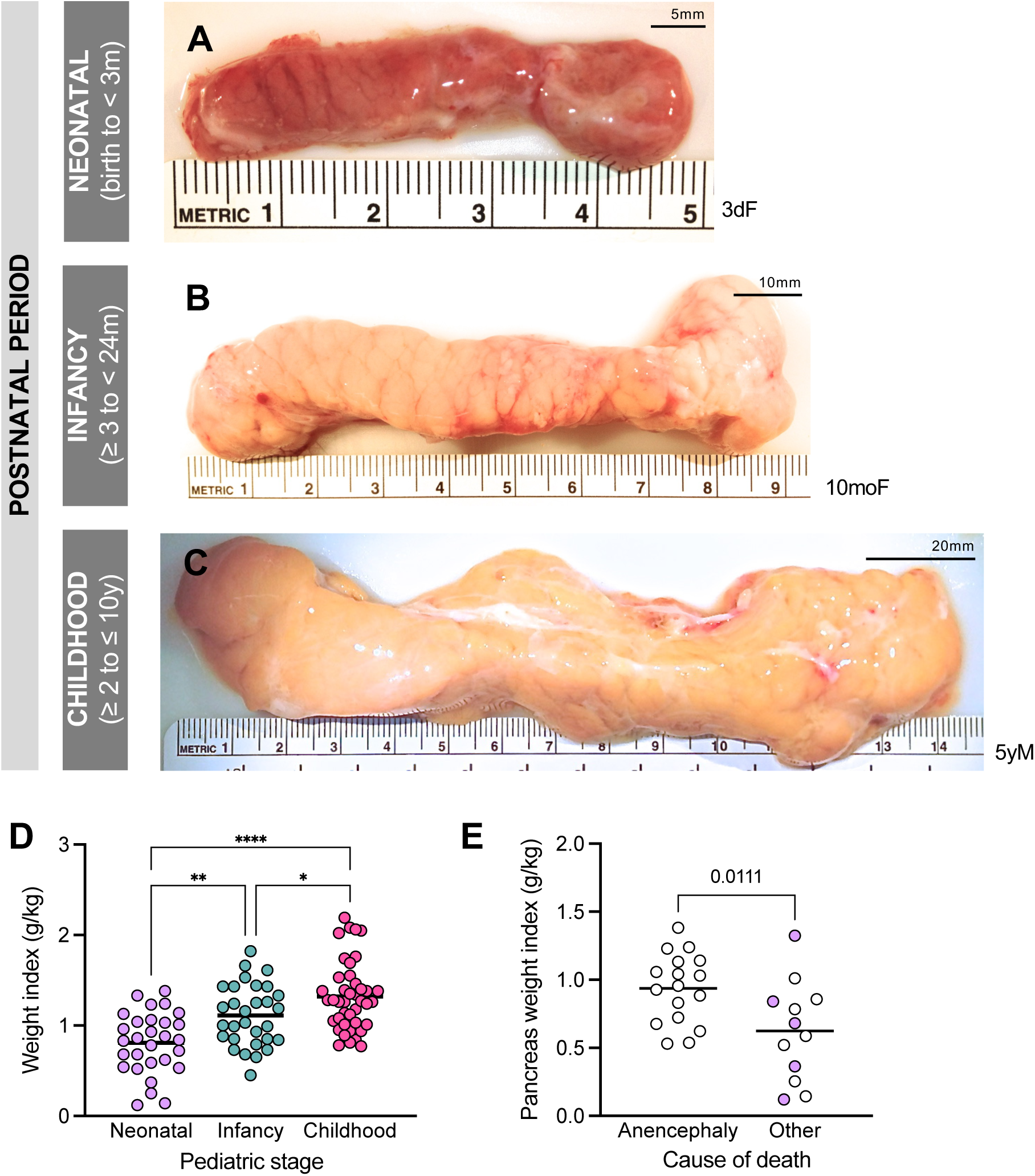
Pancreas size and pancreatic weight index increased with age and were decoupled from neural tube defect (anencephaly). **(A–C)** Images showing gross morphology and length of select postnatal pancreata from developmental stages neonatal (**A**), infancy (**B**), and childhood (**C**). **(D)** Weight index (ratio of pancreas weight to body weight) significantly increased during postnatal development (one-way ANOVA, F=18.63, *p*<0.001); however, standard deviations are not significantly different among groups (Brown-Forsythe test, F=0.06396, *p*=0.9381). See also **Figure 1, B–D. (E)** Weight index of neonatal donors stratified by cause of death. Donors aged 0-8 days denoted by white circles.

**Figure S2.**
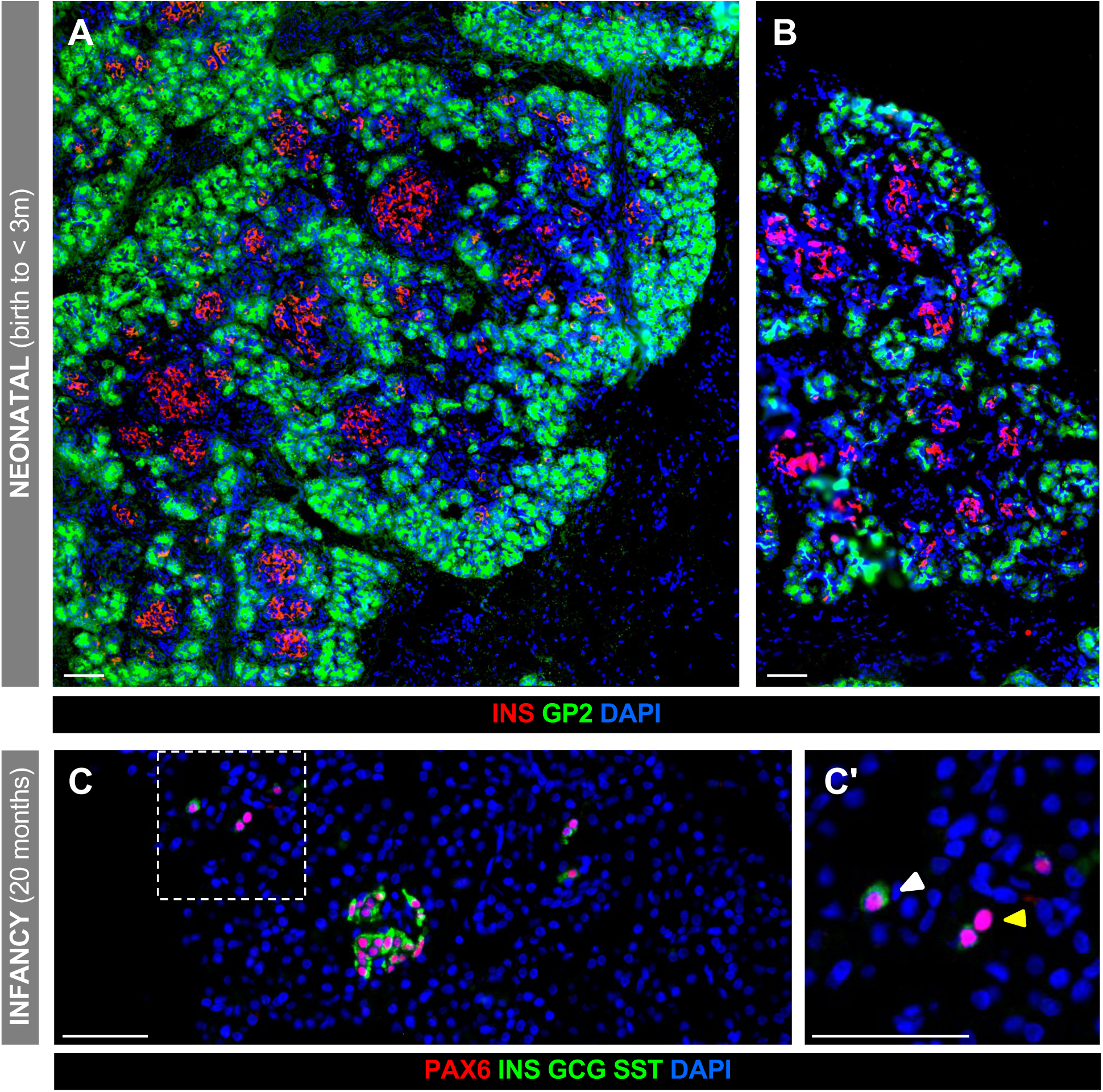
GP2+ multipotent progenitors, as well as proto-endocrine PAX6+ post-mitotic cells, persisted in human pancreas after birth. **(A–B)** Immunohistochemical staining of pancreatic sections from human neonatal donors (1 day) born at 33 (**A**) and 39.9 weeks of gestation (**B**). GP2 expression was highest around periphery of pancreatic lobes. **(C–C’)** Newly formed endocrine cells appeared around lobe periphery during the neonatal and infancy stages. Area within dashed line in **C** is magnified in **C’**; white arrowheads indicate PAX6+, hormone-positive cells; yellow arrowheads indicate PAX6+, hormone-negative cells. All scale bars are 50 μm.

**Figure S3.**
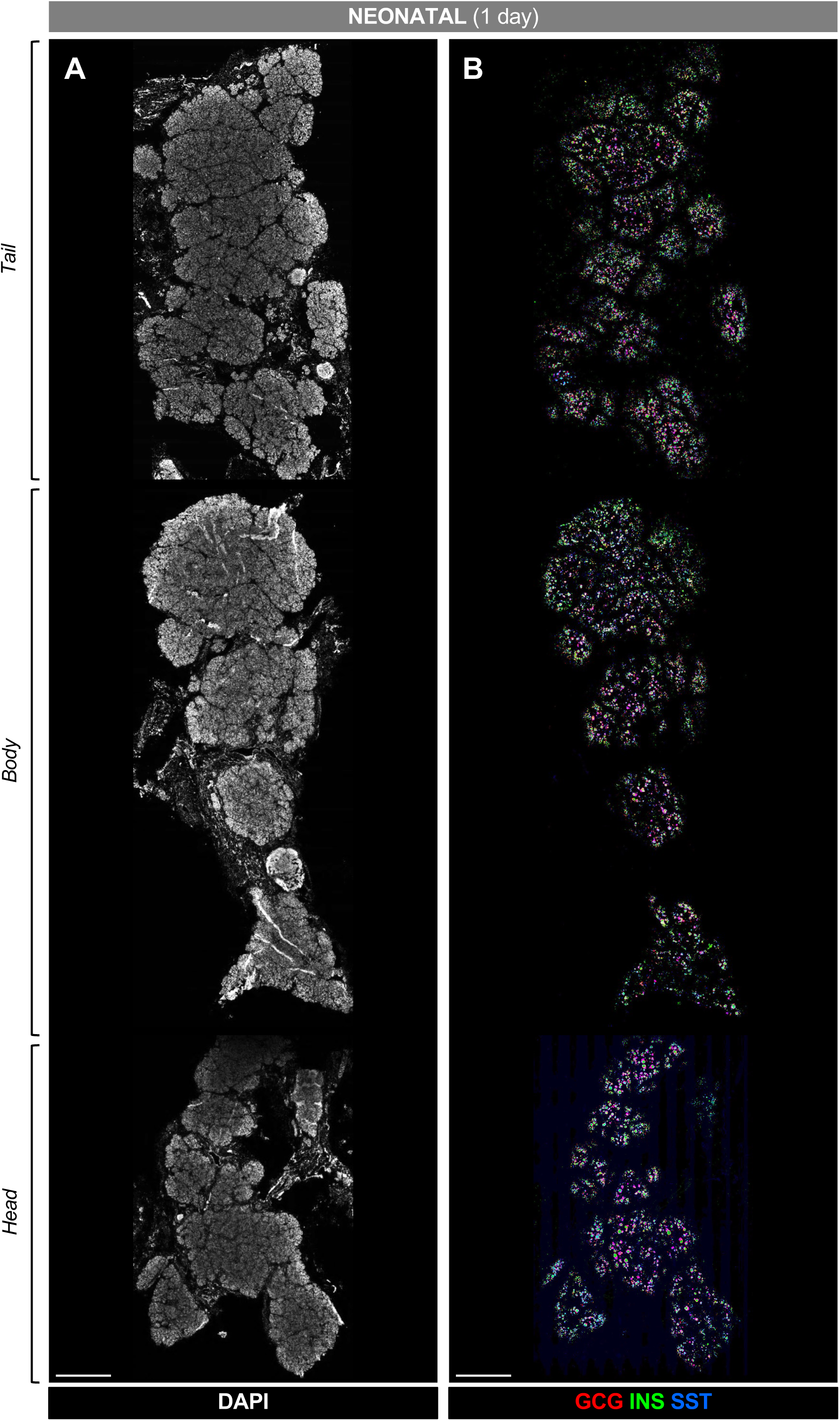
Endocrine cells were distributed throughout all pancreas regions at birth. Macro view of neonatal pancreas (aged 1 day) was generated by recombining longitudinal sections from tail, body, and head regions. Sections were counterstained with DAPI (**A**) and endocrine cells visualized by labeling for insulin (INS, green), glucagon (GCG, red), and somatostatin (SST, blue) (**B**). Scale bars, 2 mm.

**Figure S4.**
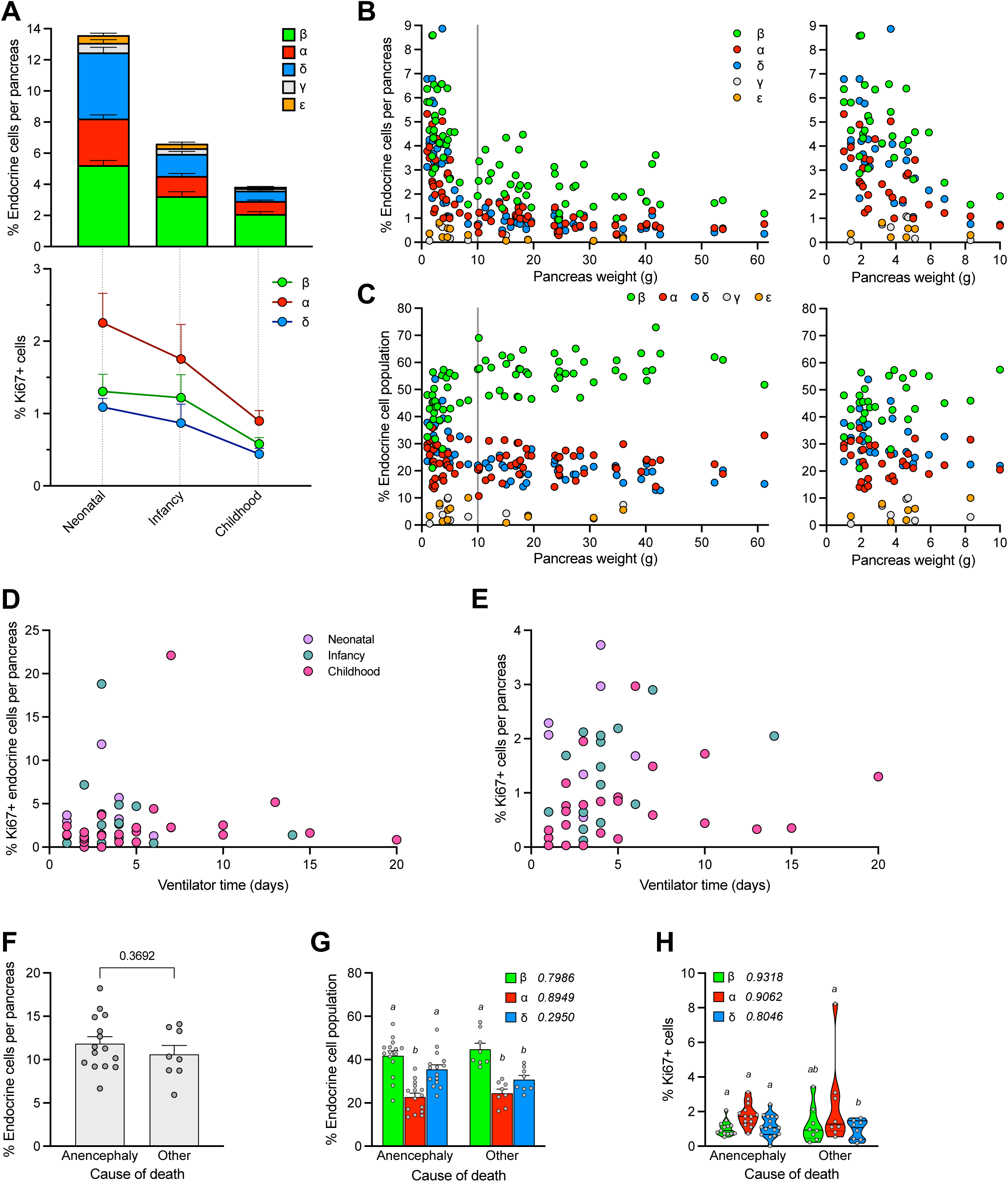
Relative proportions of β and δ cell populations changed during postnatal pancreas growth, but endocrine cell abundance, proportion, and proliferation rates were unchanged by cause of death or ventilator time prior to organ donation. **(A)** Top: number of endocrine cells per pancreas and proportion of individual endocrine cell types; overall bar height represents total density of endocrine cells (of total pancreatic cells) and fractions within bars correspond to proportions (mean + SEM) of β (green), α (red), δ (blue), ε (magenta), and γ (yellow) cells. Bottom: proliferation rates of α, β, and δ cells. See also **Figure 3D-E**. **(B–C)** Endocrine cell density (**B**) and composition (**C**) of β (green), α (red), δ (blue), ε (magenta), and γ (yellow) cells expressed in relationship to pancreas weight; α, β, δ cells, *n*=79; ε, γ cells, *n*=12. Data between y-axis and vertical line x=10 are enlarged in panels on right. **(D–E)** Neither total pancreatic (**D**) nor endocrine cell (**E**) proliferation rates varied according to ventilator time (Pearson correlation, *p*>0.05) for any individual age category (*n*=7 neonatal, *n*=16 infancy, and *n*=24 childhood donors) or combined. Ventilator time was not available for all donors (see **Table S1**). **(F–H)** Endocrine cell characteristics in study cohort did not vary by cause of death. Quantification of endocrine cell density (**F**), islet composition (**G**) and endocrine cell proliferation (**H**) of neonatal donors stratified by cause of death; *n*=15 anencephaly and *n*=9 other. Data are plotted as mean + SEM (**A**, **F**–**G**) or median ± quartiles (**H**), with each circle representing one individual. Means were analyzed by unpaired t-test (**F**) or two-way ANOVA followed by Tukey’s multiple comparisons tests (**G**–**H**). Italicized letters in **G**–**H** denote comparison among cell types within each group; bars with like letters do not statistically differ. No cell-type differences were detected between groups (*p*>0.05); *p*-values are listed next to key labels.

**Figure S5.**
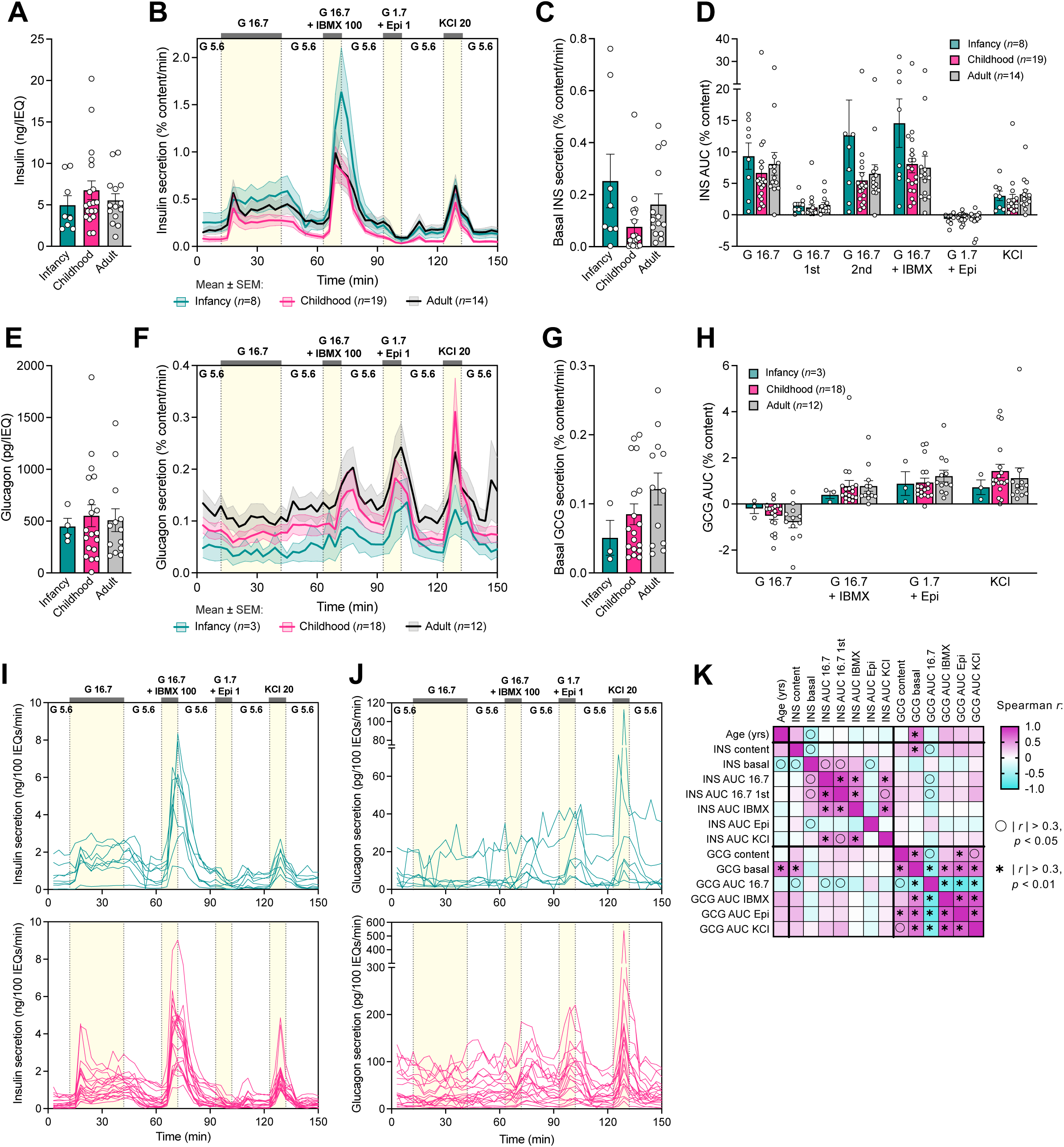
Normalization to islet hormone content masked functional changes observed in insulin and glucagon secretory responses by IEQ. **(A–D)** Insulin content (**A**) and dynamic insulin secretory response (**B**–**D**) of islets from infancy (*n*=8), childhood (*n*=19), and adult (*n*=14) independent donors measured by perifusion, normalized to insulin content. Differences in insulin secretion trajectories normalized to insulin content were analyzed by mixed-effect models (global *p*= 4.898e-16; pairwise comparisons infancy vs. childhood, *p*=4.188e-12; infancy vs. adult, *p*=9.689e-10; childhood vs. adult, *p*=9.380e-14). **(E–H)** Glucagon content (**E**) and dynamic glucagon secretory response (**F–H**) of islets from infancy (*n*=3), childhood (*n*=18), and adult (*n*=12) independent donors measured by perifusion, normalized to glucagon content. Differences in glucagon secretion trajectories normalized to glucagon content were analyzed by mixed-effect models (global *p*=1.570e-05; pairwise comparisons infancy vs. childhood, *p*=1.859e-03; infancy vs. adult, *p*=1.997e-04; childhood vs. adult, *p*=1.068e-03). Samples were excluded if glucagon fell below the limit of detection in islet extracts, and some historic samples had to be omitted due to inadequate archived material. See **Methods** for more details. Panels **D** and **H** show secretory response analyzed as area under the curve (AUC). Treatments (and concentrations) are shown along the top of the graph: Epi, epinephrine (μM); G, glucose (mM); IBMX, isobutylmethylxanthine (μM); KCl, potassium chloride (mM). Means were analyzed by one-way (**A, C, E, G**) or two-way (**D**, **H**) ANOVA followed by Tukey’s multiple comparisons tests. (**I**–**J**) IEQ-normalized insulin (**I**) and glucagon (**J**) secretory responses of islets from individual infancy (teal, top) and childhood (pink, bottom) donors; aggregated data shown in Figure 4E and **4H**. **(K)** Matrix showing Spearman correlation of IEQ-normalized secretory traits from *n*=28 islet preparations from pediatric donors. Symbols denote absolute values of Spearman *r* >0.3; ○, approximate *p*<0.05; ∗, approximate *p*<0.01.

**Figure S6.**
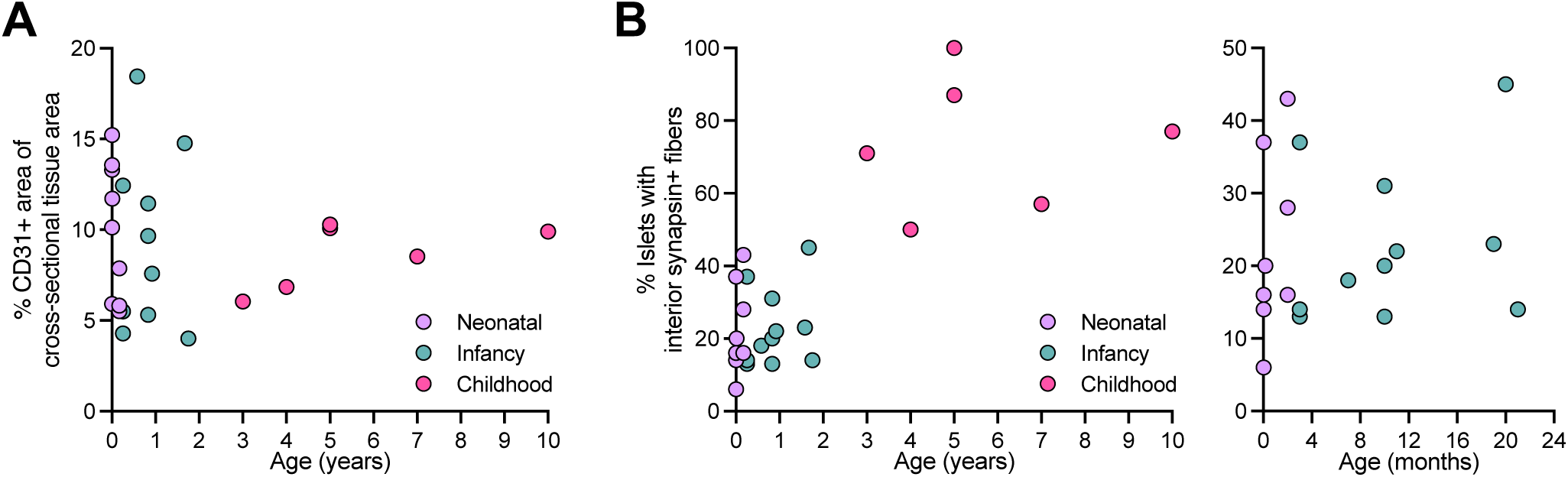
Islet innervation was delayed compared to islet vascularization. Islet vascular density (**A**; defined as percentage CD31+ area of total cross-sectional tissue area) and innervation (**B**; defined as percentage of islets with interior synapsin-1/2+ fibers) depicted across stages of postnatal pancreas development. See also Figure 5.

## SUPPLEMENTAL TABLES

**Table S1.** Donor demographic information.

**Table S2.** Primary antibodies used for immunohistochemistry.

**Table S3.** Secondary antibodies used for immunohistochemistry.

**Table S4.** Hormone assays used for assessing islet function.

**Table S5.** Comparison of dynamic perifusion profile between infancy, childhood, and adult samples

## MATERIALS AND METHODS

### Human postnatal pancreas procurement and processing

Pancreata from nondiabetic postnatal donors (**Figure S1A-C**) were obtained through partnerships with the Network of Pancreatic Organ Donors with Diabetes (nPOD), the International Institute for Advancement of Medicine (IIAM), National Disease Research Interchange (NDRI), Integrated Islet Distribution Program (IIDP). The Vanderbilt University Institutional Review Board declared that study on de-identified human pancreatic specimens does not qualify as human subject research. Donor demographic information is summarized in **Table S1**. Pancreata were received within 18-24 hours from cold clamp and maintained in cold preservation solution on ice until processing.

Upon receipt, pancreas was cleaned from connective tissue and fat, measured, and weighed, and then cross-sectional slices of pancreas with 2-3 mm thickness were obtained from the head, body and distal tail (**Figure 1A**). For donors ≥1 year of age, pancreatic slices were further divided into four quadrants. Moving from head to tail, sequential tissue slices were processed by alternating protocols: light fixation and cryopreservation, formalin fixation and paraffin embedding, fixation for electron microscopy, and snap freezing with liquid nitrogen; methods are detailed at https://dx.doi.org/10.17504/protocols.io.eq2lyp5nelx9/v2. Briefly, samples for cryopreservation were fixed in 0.1 M phosphate buffered saline (PBS) containing 4% paraformaldehyde (Electron Microscopy Sciences) for 2-3 hours on ice with mild agitation, washed in four changes of 0.1 M PBS over 2 hours, equilibrated in 30% sucrose/0.01 M PBS overnight and embedded in Tissue-Plus optimal cutting temperature (OCT) compound (Fisher Scientific). Samples for paraffin embedding were fixed in 10% neutral buffered formalin (Electron Microscopy Sciences) overnight at 4°C under mild agitation, washed in four changes of 0.1 M PBS over 3 hours, and stored in 70% ethanol at 4°C to process for paraffin embedding. Samples for electron microscopy were manually minced in fixative (2% paraformaldehyde, 2% glutaraldehyde in 0.1 M cacodylate buffer) warmed to room temperature, then incubated in fixative for 24-72 hours at 4°C under mild agitation. In certain cases, a modified procedure was used that enabled islet isolation and tissue collection from the same organ^20,85–87^.

#### Islet functional analysis

We utilized an islet phenotyping pipeline that has been applied across different disease states and to more than 500 nondiabetic adult islet preparations^7^. Islets from infancy (<2 years for functional studies, *n*=8; single 2 month old neonatal donor included with ≥3 months to <24 months infant donors), childhood (2-10 years, *n*=20), and adult (24-59 years, *n*=14) nondiabetic donors were studied in a dynamic cell perifusion system at a perifusate flow rate of 1 mL/min^86^ .

The effluent was collected at 3-minute intervals using an automatic fraction collector, then islets were retrieved and lysed with acid-ethanol solution to extract. Insulin and/or glucagon concentrations in each perifusion fraction, as well as total hormone content, were measured by radioimmunoassay (RIA) (human insulin, Millipore RI-13K; glucagon, Millipore GL-32K), enzyme-linked immunosorbent assay (ELISA) (human insulin, Mercodia 10-1132-01; glucagon, Mercodia 10-1281-01 and R&D DGCG0), or homogeneous time-resolved fluorescence (HTRF) assay (glucagon, Cisbio 62CGLPEH). Specific assays used are detailed in **Table S4**. Exclusion criteria were as follows: content-normalized traces were excluded if glucagon fell below the limit of detection in islet extracts or in the case of historic samples, if there was inadequate archived material to perform a more sensitive assay, as well as if part of the islet pellet was lost during retrieval. One content-normalized glucagon trace from the childhood group was excluded by an outlier test. Adult IEQ- or content-normalized traces were excluded if they fell outside two standard deviations of average traces obtained by over 500 adult nondiabetic islet preparations by the Human Islet Phenotyping Program (HIPP) of the Integrated Islet Distribution Program (IIDP)^7^. Area under the curve (AUC) above baseline hormone release was calculated with the trapezoidal method in GraphPad Prism 8-10 as previously described^88^, using the following time point boundaries: insulin 16.7 mM glucose, 9-60 min (1st phase 9-24 min); insulin 16.7 mM glucose + IBMX, 63-90 min; insulin 1.7 mM glucose + epinephrine, 90-114 min; insulin KCl, 120-150 min; glucagon 16.7 mM glucose, 12-51 min; glucagon 16.7 mM glucose + IBMX, 69-90 min; glucagon 1.7 mM glucose + epinephrine, 93-117 min; glucagon KCl, 120-144 min. In addition, hormone secretory trajectories were analyzed across different age groups – infancy, childhood, and adulthood – using linear mixed-effect models^55^ that take into consideration the underlying temporal correlation. In each model, restricted cubic splines were used to capture the nonlinear relationships between hormone secretory trajectories and time. The statistical significance (*p*<0.05) of the interaction between the splines and group indicators, assessed through a likelihood ratio test, confirmed differences in hormone secretory trajectories among the three groups. A similar model was employed to compare hormone secretory trajectories between infancy and either childhood or adulthood, and between childhood and adulthood. The statistical significance of the interaction between the splines and a binary group indicator confirmed these differences.

#### Immunohistochemical analysis

### Staining and imaging

Immunohistochemical analysis of pancreas was performed on 5 to 8 μm cryosections from multiple blocks from head, body and tail regions as described previously^89^.

Immunohistochemical analysis of isolated islets was performed on 10 µm cryosections of islets immobilized in type 1 collagen gels as described previously^90^ and in accordance with the Human Islet Phenotyping Program pipeline (https://dx.doi.org/10.17504/protocols.io.bqunmwve, https://dx.doi.org/10.17504/protocols.io.b2nfqdbn). For whole mount labelling, tissue was first washed in 1 mL of PBSBT containing 1X PBS + 2% BSA + 0.5% TritonX-100 for 60 minutes at 4°C with gentle rocking on a nutator and then transferred into 1 mL of fresh PBSBT for overnight incubation at 4°C. Primary and secondary antibody mixtures were made in PBSBT and added in 500 μL/tube for overnight incubation at 4°C with gentle rocking. Tissues were rinsed twice and then washed 3 times (>1 hour/each wash) in 1 mL of PBSBT at 4°C prior to addition of secondary antibody mixture and after the secondary antibody incubation. Following the immunofluorescence labeling, tissues were dehydrated in three steps at 4°C using a series of methanol solutions including 50% MeOH (30 minutes), 80% MeOH (30 minutes) and 100% MeOH (2x 30 minutes) with 1 mL/each step and stored in dark at 4°C. When ready for imaging, tissue was cleared in BABB reagent containing benzyl alcohol (Sigma A305197) and benzyl benzoate (Sigma B6630) mixed at 1:2 ratio and then kept in BABB during imaging. After imaging, tissue was transferred to 100% MeOH for storage. Primary antibodies to all antigens and their working dilutions are listed in **Table S2**. Digital images were acquired with a Zeiss LSM510 META laser scanning confocal microscope (Carl Zeiss), ScanScope FL (Leica/Aperio), and DMI6000B fluorescence microscope equipped with a DFC360FX digital camera (Leica).

### Endocrine cell abundance, proliferation, and spatial arrangement during postnatal development

Scanned images were inspected for quality and annotated by hand to exclude out-of-focus regions. Hormone and Ki67 positivity were quantified using a tuned cytonuclear v1.2 algorithm (Aperio). Endocrine cell mass was quantified by using pancreas weight and the ratio of hormone positive cells as identified by cytonuclear logarithm within the entire pancreatic section from multiple blocks representing the head, body, and tail regions. The following analysis metrics represent mean ± standard error: endocrine cell density and composition (**Figure 3D-E**, **S4A-C**) 35,844 ± 3,163 islet cells/donor and 629,419 ± 44,456 total cells/donor; endocrine cell proliferation (**Figure 3F, S4D-E**) 44,497 ± 4,027 cells/donor; islet cell arrangement (**Figure 4A-D**) 30 islets/donor, 3,050 ± 532 cells/donor.

### Postnatal endothelial cell abundance and islet innervation

Scanned images were inspected for quality and annotated by hand to exclude out-of-focus regions. Positivity for CD31 (endothelial cell marker) was quantified via tissue classifier using HALO imaging analysis software (Indica Labs). For innervation, islets (estimated diameter ≥50 μm) were annotated based on DAPI and hormone channels. The TUBB3 (neuronal marker) channel was then turned on and each islet was evaluated for whether there was TUBB3 signal ≥ 2 cell layers from the periphery of the islet (considered “interior nerve fibers”). The following analysis metrics represent mean ± standard error: endothelial cell area (**Figure 7L, S8L**) (*n*=25; 0.3004 ± 0.0459 mm^2^), islet innervation (**Figure 7P, S8M**) (*n*=25; 48 ± 7 islets/donor).

#### Statistical analysis

Statistical analysis was performed using GraphPad Prism software and R Studio. Endocrine cell populations in postnatal pancreas were compared by one- or two-way ANOVA followed by Tukey’s multiple comparisons test. Data are expressed as mean ± standard error of mean unless otherwise indicated. Two-tailed Student’s *t* test was used for analysis of statistical significance for two-group comparisons. A *p*-value less than 0.05 was considered significant. To capture the nonlinear trend of pancreas weight or weight index over age, we employed restricted cubic splines of the age variable as the primary explanatory variables in our linear regression models. The statistical significance (p value < 0.05) of the spline terms was tested by comparing the model with restricted cubic spline terms to the model without them using an F-test. Statistical details of experiments are provided in figure legends and/or Results text as appropriate.

## REFERENCES

1. Kassem, S.A., Ariel, I., Thornton, P.S., Scheimberg, I., and Glaser, B. (2000). Beta-cell proliferation and apoptosis in the developing normal human pancreas and in hyperinsulinism of infancy. Diabetes 49, 1325–1333. 10.2337/diabetes.49.8.1325.

2. Cnop, M., Hughes, S.J., Igoillo-Esteve, M., Hoppa, M.B., Sayyed, F., Laar, L. van de, Gunter, J.H., Koning, E.J.P. de, Walls, G.V., Gray, D.W.G., et al. (2009). The long lifespan and low turnover of human islet beta cells estimated by mathematical modelling of lipofuscin accumulation. Diabetologia 53, 321. 10.1007/s00125-009-1562-x.

3. Meier, J.J., Köhler, C.U., Alkhatib, B., Sergi, C., Junker, T., Klein, H.H., Schmidt, W.E., and Fritsch, H. (2010). β-cell development and turnover during prenatal life in humans. Eur J Endocrinol 162, 559–568. 10.1530/eje-09-1053.

4. Perl, S., Kushner, J.A., Buchholz, B.A., Meeker, A.K., Stein, G.M., Hsieh, M., Kirby, M., Pechhold, S., Liu, E.H., Harlan, D.M., et al. (2010). Significant human beta-cell turnover is limited to the first three decades of life as determined by in vivo thymidine analog incorporation and radiocarbon dating. J Clin Endocrinol Metabolism 95, E234–9. 10.1210/jc.2010-0932.

5. Gregg, B.E., Moore, P.C., Demozay, D., Hall, B.A., Li, M., Husain, A., Wright, A.J., Atkinson, M.A., and Rhodes, C.J. (2012). Formation of a Human β-Cell Population within Pancreatic Islets Is Set Early in Life. J Clin Endocrinol Metabolism 97, 3197–3206. 10.1210/jc.2012-1206.

6. Saisho, Y., Butler, A.E., Manesso, E., Elashoff, D., Rizza, R.A., and Butler, P.C. (2012). β-cell mass and turnover in humans: effects of obesity and aging. Diabetes Care 36, 111–117. 10.2337/dc12-0421.

7. Evans-Molina, C., Pettway, Y.D., Saunders, D.C., Sharp, S.A., Bate, T.SR., Sun, H., Durai, H., Mei, S., Coldren, A., Davis, C., et al. (2024). Heterogeneous endocrine cell composition defines human islet functional phenotypes. bioRxiv, 2024.11.20.623809. 10.1101/2024.11.20.623809.

8. Arda, H.E., Li, L., Tsai, J., Torre, E.A., Rosli, Y., Peiris, H., Spitale, R.C., Dai, C., Gu, X., Qu, K., et al. (2016). Age-Dependent Pancreatic Gene Regulation Reveals Mechanisms Governing Human β Cell Function. Cell Metab 23, 909–920. 10.1016/j.cmet.2016.04.002.

9. Ziegler, A.G., Rewers, M., Simell, O., Simell, T., Lempainen, J., Steck, A., Winkler, C., Ilonen, J., Veijola, R., Knip, M., et al. (2013). Seroconversion to Multiple Islet Autoantibodies and Risk of Progression to Diabetes in Children. JAMA 309, 2473–2479. 10.1001/jama.2013.6285.

10. Warncke, K., Weiss, A., Achenbach, P., Berge, T. von dem, Berner, R., Casteels, K., Groele, L., Hatzikotoulas, K., Hommel, A., Kordonouri, O., et al. (2022). Elevations in blood glucose before and after the appearance of islet autoantibodies in children. J Clin Invest 132, e162123. 10.1172/jci162123.

11. Castaing, M., Duvillié, B., Quemeneur, E., Basmaciogullari, A., and Scharfmann, R. (2005). Ex vivo analysis of acinar and endocrine cell development in the human embryonic pancreas. Dev Dynam 234, 339–345. 10.1002/dvdy.20547.

12. Li, Z., Tremmel, D.M., Ma, F., Yu, Q., Ma, M., Delafield, D.G., Shi, Y., Wang, B., Mitchell, S.A., Feeney, A.K., et al. (2021). Proteome-wide and matrisome-specific alterations during human pancreas development and maturation. Nat Commun 12, 1020. 10.1038/s41467-021-21261-w.

13. Jeon, J., Correa-Medina, M., Ricordi, C., Edlund, H., and Diez, J.A. (2009). Endocrine cell clustering during human pancreas development. J Histochem Cytochem 57, 811–824. 10.1369/jhc.2009.953307.

14. Jennings, R.E., Berry, A.A., Kirkwood-Wilson, R., Roberts, N.A., Hearn, T., Salisbury, R.J., Blaylock, J., Hanley, K.P., and Hanley, N.A. (2013). Development of the Human Pancreas From Foregut to Endocrine Commitment. Diabetes 62, 3514–3522. 10.2337/db12-1479.

15. Piper, K., Brickwood, S., Turnpenny, L.W., Cameron, I.T., Ball, S.G., Wilson, D.I., and Hanley, N.A. (2004). Beta cell differentiation during early human pancreas development. J Endocrinol 181, 11–23. 10.1677/joe.0.1810011.

16. Lyttle, B.M., Li, J., Krishnamurthy, M., Fellows, F., Wheeler, M.B., Goodyer, C.G., and Wang, R. (2008). Transcription factor expression in the developing human fetal endocrine pancreas. Diabetologia 51, 1169–1180. 10.1007/s00125-008-1006-z.

17. Villalba, A., Gitton, Y., Inoue, M., Aiello, V., Blain, R., Toupin, M., Mazaud-Guittot, S., Rachdi, L., Semb, H., Chédotal, A., et al. (2024). A 3D atlas of the human developing pancreas to explore progenitor proliferation and differentiation. Diabetologia 67, 1066–1078. 10.1007/s00125-024-06143-2.

18. Pauerstein, P.T., Sugiyama, T., Stanley, S.E., McLean, G.W., Wang, J., Martín, M.G., and Kim, S.K. (2015). Dissecting Human Gene Functions Regulating Islet Development With Targeted Gene Transduction. Diabetes 64, 3037–3049. 10.2337/db15-0042.

19. Bouwens, L., Lu, W.G., and Krijger, R.D. (1997). Proliferation and differentiation in the human fetal endocrine pancreas. Diabetologia 40, 398–404. 10.1007/s001250050693.

20. Dai, C., Hang, Y., Shostak, A., Poffenberger, G., Hart, N., Prasad, N., Phillips, N., Levy, S.E., Greiner, D.L., Shultz, L.D., et al. (2017). Age-dependent human β cell proliferation induced by glucagon-like peptide 1 and calcineurin signaling. J Clin Invest 127, 3835–3844. 10.1172/jci91761.

21. Banaei-Bouchareb, L., Peuchmaur, M., Czernichow, P., and Polak, M. (2006). A transient microenvironment loaded mainly with macrophages in the early developing human pancreas. J Endocrinol 188, 467–480. 10.1677/joe.1.06225.

22. Amella, C., Cappello, F., Kahl, P., Fritsch, H., Lozanoff, S., and Sergi, C. (2008). Spatial and temporal dynamics of innervation during the development of fetal human pancreas. Neuroscience 154, 1477–1487. 10.1016/j.neuroscience.2008.04.050.

23. Gorczyca, J., Litwin, J.A., Pitynski, K., and Miodonski, A.J. (2010). Vascular system of human fetal pancreas demonstrated by corrosion casting and scanning electron microscopy. Anat Sci Int 85, 235–240. 10.1007/s12565-010-0084-4.

24. Gorczyca, J., Tomaszewski, K.A., Henry, B.M., Pękala, P.A., Pasternak, A., Mizia, E., and Walocha, J.A. (2017). The Vascular Microarchitecture of the Human Fetal Pancreas: A Corrosion Casting and Scanning Electron Microscopy Study. Pancreas 46, 124–130. 10.1097/mpa.0000000000000719.

25. Krivova, Y.S., Proshchina, A.E., Otlyga, D.A., Leonova, O.G., and Saveliev, S.V. (2022). Prenatal development of sympathetic innervation of the human pancreas. Ann Anat 240, 151880. 10.1016/j.aanat.2021.151880.

26. Nekrep, N., Wang, J., Miyatsuka, T., and German, M.S. (2008). Signals from the neural crest regulate beta-cell mass in the pancreas. Development 135, 2151–2160. 10.1242/dev.015859.

27. Borden, P., Houtz, J., Leach, S.D., and Kuruvilla, R. (2013). Sympathetic Innervation during Development Is Necessary for Pancreatic Islet Architecture and Functional Maturation. Cell Rep 4, 287–301. 10.1016/j.celrep.2013.06.019.

28. Reinert, R.B., Cai, Q., Hong, J.-Y., Plank, J.L., Aamodt, K., Prasad, N., Aramandla, R., Dai, C., Levy, S.E., Pozzi, A., et al. (2014). Vascular endothelial growth factor coordinates islet innervation via vascular scaffolding. Development 141, 1480–1491. 10.1242/dev.098657.

29. Lee, I. (2016). Human pancreatic islets develop through fusion of distinct β and α/δ islets. Dev Growth Differ 58, 635–640. 10.1111/dgd.12308.

30. Proshchina, A.E., Krivova, Y.S., Barabanov, V.M., and Saveliev, S.V. (2019). Pancreatic endocrine cell arrangement during human ontogeny. Acta Histochem 121, 638–645. 10.1016/j.acthis.2019.05.010.

31. Bader, E., Migliorini, A., Gegg, M., Moruzzi, N., Gerdes, J., Roscioni, S.S., Bakhti, M., Brandl, E., Irmler, M., Beckers, J., et al. (2016). Identification of proliferative and mature β-cells in the islets of Langerhans. Nature 535, 430–434. 10.1038/nature18624.

32. Avrahami, D., Li, C., Zhang, J., Schug, J., Avrahami, R., Rao, S., Stadler, M.B., Burger, L., Schübeler, D., Glaser, B., et al. (2015). Aging-Dependent Demethylation of Regulatory Elements Correlates with Chromatin State and Improved β Cell Function. Cell Metab 22, 619– 632. 10.1016/j.cmet.2015.07.025.

33. Dhawan, S., Tschen, S.-I., Zeng, C., Guo, T., Hebrok, M., Matveyenko, A., and Bhushan, A. (2015). DNA methylation directs functional maturation of pancreatic β cells. J Clin Invest 125, 2851–2860. 10.1172/jci79956.

34. Xu, J., Murphy, S.L., Kochanek, K.D., and Arias, E. (2022). Mortality in the United States, 2021. NCHS Data Brief, no 456, 1–8. 10.15620/cdc:122516.

35. Goyal, N. (2020). The newborn infant. In Nelson Textbook of Pediatrics, R. M. Kliegman, J. W. S. G. III, N. J. Blum, S. S. Shah, R. C. Tasker, and K. M. Wilson, eds. (Elsevier).

36. Ellis, R. (1951). Phases of Postnatal Growth. Brit J Nutr 5, 151–157. 10.1079/bjn19510018.

37. Hill, M.A. (2020). Embryology. Embryology iBooks. https://embryology.med.unsw.edu.au/embryology/index.php/Embryology_iBooks.

38. Saunders, D.C., Messmer, J., Kusmartseva, I., Beery, M.L., Yang, M., Atkinson, M.A., Powers, A.C., Cartailler, J.-P., and Brissova, M. (2020). Pancreatlas: Applying an Adaptable Framework to Map the Human Pancreas in Health and Disease. Patterns 1, 100120. 10.1016/j.patter.2020.100120.

39. Virostko, J., Williams, J., Hilmes, M., Bowman, C., Wright, J.J., Du, L., Kang, H., Russell, W.E., Powers, A.C., and Moore, D.J. (2019). Pancreas Volume Declines During the First Year After Diagnosis of Type 1 Diabetes and Exhibits Altered Diffusion at Disease Onset. Diabetes Care 42, 248–257. 10.2337/dc18-1507.

40. Campbell-Thompson, M.L., Filipp, S.L., Grajo, J.R., Nambam, B., Beegle, R., Middlebrooks, E.H., Gurka, M.J., Atkinson, M.A., Schatz, D.A., and Haller, M.J. (2018). Relative Pancreas Volume Is Reduced in First-Degree Relatives of Patients With Type 1 Diabetes. Diabetes Care 42, 281–287. 10.2337/dc18-1512.

41. Cogger, K.F., Sinha, A., Sarangi, F., McGaugh, E.C., Saunders, D., Dorrell, C., Mejia-Guerrero, S., Aghazadeh, Y., Rourke, J.L., Screaton, R.A., et al. (2017). Glycoprotein 2 is a specific cell surface marker of human pancreatic progenitors. Nat Commun 8, 331. 10.1038/s41467-017-00561-0.

42. Aghazadeh, Y., Sarangi, F., Poon, F., Nkennor, B., McGaugh, E.C., Nunes, S.S., and Nostro, M.C. (2022). GP2-enriched pancreatic progenitors give rise to functional beta cells in vivo and eliminate the risk of teratoma formation. Stem Cell Rep 17, 964–978. 10.1016/j.stemcr.2022.03.004.

43. Schaffer, A.E., Freude, K.K., Nelson, S.B., and Sander, M. (2010). Nkx6 Transcription Factors and Ptf1a Function as Antagonistic Lineage Determinants in Multipotent Pancreatic Progenitors. Dev. Cell 18, 1022–1029. 10.1016/j.devcel.2010.05.015.

44. Benitez, C.M., Goodyer, W.R., and Kim, S.K. (2012). Deconstructing Pancreas Developmental Biology. Csh Perspect Biol 4, a012401. 10.1101/cshperspect.a012401.

45. Masui, T., Swift, G.H., Hale, M.A., Meredith, D.M., Johnson, J.E., and MacDonald, R.J. (2008). Transcriptional Autoregulation Controls Pancreatic Ptf1a Expression during Development and Adulthood. Mol. Cell. Biol. 28, 5458–5468. 10.1128/mcb.00549-08.

46. Pan, F.C., and Wright, C. (2011). Pancreas organogenesis: from bud to plexus to gland. Dev Dynam 240, 530–565. 10.1002/dvdy.22584.

47. Glorieux, L., Sapala, A., Willnow, D., Moulis, M., Salowka, A., Darrigrand, J.-F., Edri, S., Schonblum, A., Sakhneny, L., Schaumann, L., et al. (2022). Development of a 3D atlas of the embryonic pancreas for topological and quantitative analysis of heterologous cell interactions. Development 149, dev199655. 10.1242/dev.199655.

48. Smeets, S., Stangé, G., Leuckx, G., Roelants, L., Cools, W., Paep, D.L.D., Ling, Z., Leu, N.D., and Veld, P. in’t (2019). Evidence of Tissue Repair in Human Donor Pancreas After Prolonged Duration of Stay in Intensive Care. Diabetes 69, 401–412. 10.2337/db19-0529.

49. Blaas, H. -G. K., and Eik-Nes, S.H. (2009). Sonoembryology and early prenatal diagnosis of neural anomalies. Prenat Diagn 29, 312–325. 10.1002/pd.2170.

50. Copp, A.J., and Greene, N.D.E. (2013). Neural tube defects—disorders of neurulation and related embryonic processes. Wiley Interdiscip Rev Dev Biology 2, 213–227. 10.1002/wdev.71.

51. Balamurugan, A.N., Chang, Y., Bertera, S., Sands, A., Shankar, V., Trucco, M., and Bottino, R. (2006). Suitability of human juvenile pancreatic islets for clinical use. Diabetologia 49, 1845– 1854. 10.1007/s00125-006-0318-0.

52. Brissova, M., Haliyur, R., Saunders, D., Shrestha, S., Dai, C., Blodgett, D.M., Bottino, R., Campbell-Thompson, M., Aramandla, R., Poffenberger, G., et al. (2018). α Cell Function and Gene Expression Are Compromised in Type 1 Diabetes. Cell Reports 22, 2667–2676. 10.1016/j.celrep.2018.02.032.

53. Walker, J.T., Saunders, D.C., Rai, V., Chen, H.-H., Orchard, P., Dai, C., Pettway, Y.D., Hopkirk, A.L., Reihsmann, C.V., Tao, Y., et al. (2021). Genetic risk converges on regulatory networks mediating early type 2 diabetes. Nature, 1–9. 10.1038/s41586-023-06693-2.

54. Hart, N.J., Aramandla, R., Poffenberger, G., Fayolle, C., Thames, A.H., Bautista, A., Spigelman, A.F., Babon, J.A.B., DeNicola, M.E., Dadi, P.K., et al. (2018). Cystic fibrosis–related diabetes is caused by islet loss and inflammation. Jci Insight 3, e98240. 10.1172/jci.insight.98240.

55. Fitzmaurice, G.M., Laird, N.M., and Ware, J.H. (2011). Applied longitudinal analysis. Wiley.

56. Sarkar, S.A., Kobberup, S., Wong, R., Lopez, A.D., Quayum, N., Still, T., Kutchma, A., Jensen, J.N., Gianani, R., Beattie, G.M., et al. (2008). Global gene expression profiling and histochemical analysis of the developing human fetal pancreas. Diabetologia 51, 285–297. 10.1007/s00125-007-0880-0.

57. Roost, M.S., Iperen, L. van, Bernardo, A. de M., Mummery, C.L., Carlotti, F., Koning, E.J. de, and Lopes, S.M.C. de S. (2014). Lymphangiogenesis and angiogenesis during human fetal pancreas development. Vasc Cell 6, 22. 10.1186/2045-824x-6-22.

58. Proshchina, A.E., Krivova, Y.S., Barabanov, V.M., and Saveliev, S.V. (2014). Ontogeny of Neuro-Insular Complexes and Islets Innervation in the Human Pancreas. Front Endocrinol 5, 57. 10.3389/fendo.2014.00057.

59. Richardson, T.M., Saunders, D.C., Haliyur, R., Shrestha, S., Cartailler, J.-P., Reinert, R.B., Petronglo, J., Bottino, R., Aramandla, R., Bradley, A.M., et al. (2023). Human pancreatic capillaries and nerve fibers persist in type 1 diabetes despite beta cell loss. Am J Physiol-Endoc M 324, E251–E267. 10.1152/ajpendo.00246.2022.

60. Brissova, M., Fowler, M.J., Nicholson, W.E., Chu, A., Hirshberg, B., Harlan, D.M., and Powers, A.C. (2005). Assessment of Human Pancreatic Islet Architecture and Composition by Laser Scanning Confocal Microscopy. J Histochem Cytochem 53, 1087–1097. 10.1369/jhc.5c6684.2005.

61. Drigo, R.A. e, Ali, Y., Diez, J., Srinivasan, D.K., Berggren, P.-O., and Boehm, B.O. (2015). New insights into the architecture of the islet of Langerhans: a focused cross-species assessment. Diabetologia 58, 2218–2228. 10.1007/s00125-015-3699-0.

62. Bonner-Weir, S., Sullivan, B.A., and Weir, G.C. (2015). Human Islet Morphology Revisited. J Histochem Cytochem 63, 604–612. 10.1369/0022155415570969.

63. Liu, Y., Basty, N., Whitcher, B., Bell, J.D., Sorokin, E.P., Bruggen, N. van, Thomas, E.L., and Cule, M. (2021). Genetic architecture of 11 organ traits derived from abdominal MRI using deep learning. eLife 10, e65554. 10.7554/elife.65554.

64. Rahier, J., Wallon, J., and Henquin, J.C. (1980). Abundance of somatostatin cells in the human neonatal pancreas. Diabetologia 18, 251–254. 10.1007/bf00251925.

65. Rahier, J., Wallon, J., and Henquin, J.C. (1981). Cell populations in the endocrine pancreas of human neonates and infants. Diabetologia 20, 540–546. 10.1007/bf00252762.

66. Malenczyk, K., Keimpema, E., Piscitelli, F., Calvigioni, D., Björklund, P., Mackie, K., Marzo, V.D., Hökfelt, T.G., Dobrzyn, A., and Harkany, T. (2015). Fetal endocannabinoids orchestrate the organization of pancreatic islet microarchitecture. Proc National Acad Sci 112, E6185–94. 10.1073/pnas.1519040112.

67. Masjkur, J., Poser, S.W., Nikolakopoulou, P., Chrousos, G., McKay, R.D., Bornstein, S.R., Jones, P.M., and Androutsellis-Theotokis, A. (2016). Endocrine Pancreas Development and Regeneration: Noncanonical Ideas From Neural Stem Cell Biology. Diabetes 65, 314–330. 10.2337/db15-1099.

68. Pauerstein, P.T., Tellez, K., Willmarth, K.B., Park, K., Hsueh, B., Arda, E.H., Gu, X., Aghajanian, H., Deisseroth, K., Epstein, J.A., et al. (2017). A radial axis defined by Semaphorin to Neuropilin signaling controls pancreatic islet morphogenesis. Development 144, dev.148684. 10.1242/dev.148684.

69. Scharfmann, R., Xiao, X., Heimberg, H., Mallet, J., and Ravassard, P. (2008). Beta Cells within Single Human Islets Originate from Multiple Progenitors. Plos One 3, e3559. 10.1371/journal.pone.0003559.

70. Bonfanti, P., Nobecourt, E., Oshima, M., Albagli-Curiel, O., Laurysens, V., Stangé, G., Sojoodi, M., Heremans, Y., Heimberg, H., and Scharfmann, R. (2015). Ex Vivo Expansion and Differentiation of Human and Mouse Fetal Pancreatic Progenitors Are Modulated by Epidermal Growth Factor. Stem Cells Dev 24, 1766–1778. 10.1089/scd.2014.0550.

71. Veres, A., Faust, A.L., Bushnell, H.L., Engquist, E.N., Kenty, J.H., Harb, G., Poh, Y.-C.C., Sintov, E., Gürtler, M., Pagliuca, F.W., et al. (2019). Charting cellular identity during human in vitro β-cell differentiation. Nature 569, 368–373. 10.1038/s41586-019-1168-5.

72. Lorberbaum, D.S., Kishore, S., Rosselot, C., Sarbaugh, D., Brooks, E.P., Aragon, E., Xuan, S., Simon, O., Ghosh, D., Mendelsohn, C., et al. (2020). Retinoic acid signaling within pancreatic endocrine progenitors regulates mouse and human β cell specification. Development 147, dev189977. 10.1242/dev.189977.

73. Gonçalves, C.A., Larsen, M., Jung, S., Stratmann, J., Nakamura, A., Leuschner, M., Hersemann, L., Keshara, R., Perlman, S., Lundvall, L., et al. (2021). A 3D system to model human pancreas development and its reference single-cell transcriptome atlas identify signaling pathways required for progenitor expansion. Nat Commun 12, 3144. 10.1038/s41467-021-23295-6.

74. Ameri, J., Borup, R., Prawiro, C., Ramond, C., Schachter, K.A., Scharfmann, R., and Semb, H. (2017). Efficient Generation of Glucose-Responsive Beta Cells from Isolated GP2+ Human Pancreatic Progenitors. Cell Rep 19, 36–49. 10.1016/j.celrep.2017.03.032.

75. Villani, V., Thornton, M.E., Zook, H.N., Crook, C.J., Grubbs, B.H., Orlando, G., Filippo, R.D., Ku, H.T., and Perin, L. (2019). SOX9+/PTF1A+ Cells Define the Tip Progenitor Cells of the Human Fetal Pancreas of the Second Trimester. Stem Cell Transl Med 8, 1249–126. 10.1002/sctm.19-0231.

76. Ramond, C., Glaser, N., Berthault, C., Ameri, J., Kirkegaard, J.S., Hansson, M., Honoré, C., Semb, H., and Scharfmann, R. (2017). Reconstructing human pancreatic differentiation by mapping specific cell populations during development. eLife 6, e27564. 10.7554/elife.27564.

77. O, S. de la, Liu, Z., Sun, H., Yu, S.K., Wong, D.M., Chu, E., Rao, S.A., Eng, N., Peixoto, G., Bouza, J., et al. (2022). Single-Cell Multi-Omic Roadmap of Human Fetal Pancreatic Development. Biorxiv, 2022.02.17.480942. 10.1101/2022.02.17.480942.

78. Olaniru, O.E., Kadolsky, U., Kannambath, S., Vaikkinen, H., Fung, K., Dhami, P., and Persaud, S.J. (2022). Single-cell transcriptomic and spatial landscapes of the developing human pancreas. Cell Metab 1, 184–199.e5. 10.1016/j.cmet.2022.11.009.

79. Reinert, R.B., Cai, Q., Hong, J.-Y.Y., Plank, J.L., Aamodt, K., Prasad, N., Aramandla, R., Dai, C., Levy, S.E., Pozzi, A., et al. (2014). Vascular endothelial growth factor coordinates islet innervation via vascular scaffolding. Development 141, 1480–1491. 10.1242/dev.098657.

80. Richardson, T.M., Saunders, D.C., Haliyur, R., Shrestha, S., Cartailler, J.-P., Reinert, R.B., Petronglo, J., Bottino, R., Aramandla, R., Bradley, A.M., et al. (2023). Human pancreatic capillaries and nerve fibers persist in type 1 diabetes despite beta cell loss. Am. J. Physiol.- Endocrinol. Metab. 324, E251–E267. 10.1152/ajpendo.00246.2022.

81. Bocian-Sobkowska, J., Zabel, M., Wozniak, W., and Surdyk-Zasada, J. (1999). Polyhormonal aspect of the endocrine cells of the human fetal pancreas. Histochem Cell Biol 112, 147–153. 10.1007/s004180050401.

82. Andralojc, K.M., Mercalli, A., Nowak, K.W., Albarello, L., Calcagno, R., Luzi, L., Bonifacio, E., Doglioni, C., and Piemonti, L. (2008). Ghrelin-producing epsilon cells in the developing and adult human pancreas. Diabetologia 52, 486–493. 10.1007/s00125-008-1238-y.

83. Krivova, Y.S., Barabanov, V.M., Proshchina, A.E., and Savel’ev, S.V. (2014). Distribution of chromogranin A in human fetal pancreas. Bulletin of experimental biology and medicine 156, 865–868. 10.1007/s10517-014-2471-7.

84. Polak, M., Bouchareb-Banaei, L., Scharfmann, R., and Czernichow, P. (2000). Early pattern of differentiation in the human pancreas. Diabetes 49, 225–232. 10.2337/diabetes.49.2.225.

85. Balamurugan, A.N., Chang, Y., Fung, J.J., Trucco, M., and Bottino, R. (2003). Flexible Management of Enzymatic Digestion Improves Human Islet Isolation Outcome from Sub-Optimal Donor Pancreata. Am J Transplant 3, 1135–1142. 10.1046/j.1600-6143.2003.00184.x.

86. Brissova, M., Haliyur, R., Saunders, D., Shrestha, S., Dai, C., Blodgett, D.M., Bottino, R., Campbell-Thompson, M., Aramandla, R., Poffenberger, G., et al. (2018). α Cell Function and Gene Expression Are Compromised in Type 1 Diabetes. Cell Rep 22, 2667–2676. 10.1016/j.celrep.2018.02.032.

87. Walker, J.T., Saunders, D.C., Rai, V., Chen, H.-H., Orchard, P., Dai, C., Pettway, Y.D., Hopkirk, A.L., Reihsmann, C.V., Tao, Y., et al. (2023). Genetic risk converges on regulatory networks mediating early type 2 diabetes. Nature 624, 621–629. 10.1038/s41586-023-06693-2.

88. Walker, J.T., Haliyur, R., Nelson, H.A., Ishahak, M., Poffenberger, G., Aramandla, R., Reihsmann, C., Luchsinger, J.R., Saunders, D.C., Wang, P., et al. (2020). Integrated human pseudoislet system and microfluidic platform demonstrate differences in GPCR signaling in islet cells. JCI Insight 5, e137017. 10.1172/jci.insight.137017.

89. Brissova, M., Aamodt, K., Brahmachary, P., Prasad, N., Hong, J.-Y., Dai, C., Mellati, M., Shostak, A., Poffenberger, G., Aramandla, R., et al. (2014). Islet Microenvironment, Modulated by Vascular Endothelial Growth Factor-A Signaling, Promotes β Cell Regeneration. Cell Metab 19, 498–511. 10.1016/j.cmet.2014.02.001.

90. Dai, C., Brissova, M., Hang, Y., Thompson, C., Poffenberger, G., Shostak, A., Chen, Z., Stein, R., and Powers, A.C. (2012). Islet-enriched gene expression and glucose-induced insulin secretion in human and mouse islets. Diabetologia 55, 707–718. 10.1007/s00125-011-2369-0.

