## Supplemental Table S1 for "Mapping histologic and functional maturation of human endocrine pancreas across early postnatal periods"

**Table S1. Donor characteristics, sample types, and experimental usage**

Adapted from Hart NJ, Powers AC (2018), Checklist for Reporting Human Islet Preparations Used in Research, *Diabetologia*, doi.org/10.1007/s00125-018-4772-2.

| | Donor information | | | | | | | | | | | | | | | | Assays | | | | | | | | | | |
|---|---|---|---|---|---|---|---|---|---|---|---|---|---|---|---|---|---|---|---|---|---|---|---|---|---|---|---|
| Unique identifier | Age category | Age | Sex | Race/ethnicity | BMI ($kg/m^2$) | HbA1c (ng/mL) | Cause of death | Ventilator time (days) | Source of islets/ tissue | Islet isolation center (if applicable) | Function assessed? | Est. purity (%) | Hand-picked? | Warm ischemia time (hrs) | Cold ischemia time (hrs) | Culture time (hrs) | Endo dens | α/β/δ comp | α/β/δ prolif | α/β/δ dens | Panc weight | Innervation | Vasculature | Architecture | ε/γ comp | Perifusion | Islet gels |
| HDL037 | Neonatal | 0d | M | Asian | NR | NR | Anencephaly | 0 | IIAM | | N | | | | | | • | • | • | • | • | | | | | | |
| HDL089 | Neonatal | 0d | F | Caucasian | 14.0 | NR | Anencephaly | NR | IIAM | | N | | | 2.000 | | | | | | | • | | | | | | |
| DON51 | Neonatal | 1d | M | Caucasian | NR | NR | Anencephaly | 0 | IIAM | | N | | | | | | • | • | • | • | • | | • | | | | |
| DON69 | Neonatal | 1d | M | Caucasian | NR | NR | Anencephaly | 0 | IIAM | | N | | | | | | • | • | • | • | • | | • | | | | |
| DON79 | Neonatal | 1d | F | Caucasian | 12.4 | NR | Anencephaly | 0 | IIAM | Pittsburgh | N | | | 1.450 | | | • | • | • | • | • | • | • | | | | |
| DON9 | Neonatal | 1d | F | Caucasian | 12.5 | NR | Anencephaly | 0 | IIAM | | N | | | | | | • | • | • | • | • | • | | • | | | |
| DON15 | Neonatal | 1d | F | Caucasian | 12.5 | NR | Anencephaly | 0 | IIAM | | N | | | | | | • | • | • | • | • | | | • | | | |
| HDL003 | Neonatal | 1d | F | Caucasian | 14.2 | <2.5 | Anencephaly | 0 | OPO | | N | | | | | | • | • | • | • | • | | | | | | |
| HDL022 | Neonatal | 1d | F | Caucasian | 13.0 | 5.1 | Anencephaly | 0 | IIAM | | N | | | | | | • | • | • | • | • | | | | | | |
| HDL007 | Neonatal | 1d | M | Caucasian | 13.3 | 5.6 | Anencephaly | 0 | OPO | | N | | | | | | • | • | • | • | • | | | | • | | |
| DON66 | Neonatal | 1d | F | Caucasian | 12.6 | NR | Anencephaly | 1 | IIAM | | N | | | | | | • | • | • | • | • | • | • | • | | | |
| DON112 | Neonatal | 1d | F | Hispanic or Latino | NR | NR | Anencephaly | NR | IIAM | | N | | | | | | | | • | • | | | | | | | |
| HDL059 | Neonatal | 1d | F | Unknown | NR | NR | Anencephaly | NR | IIAM | | N | | | | | | | | | | • | | | | | | |
| DON74 | Neonatal | 1d | F | Unknown | NR | NR | Anencephaly | NR | IIAM | Pittsburgh | N | | | | | | • | • | • | • | | | | | | | • |
| DON80 | Neonatal | 1d | F | Unknown | NR | NR | Anencephaly | NR | IIAM | Pittsburgh | N | | | 1.400 | | | • | • | • | • | • | • | • | | | | |
| HDL058 | Neonatal | 1d | M | Hispanic or Latino | 11.0 | NR | Anencephaly | NR | IIAM | | N | | | 1.700 | | | | | | | • | | | | | | |
| DON17 | Neonatal | 1d | M | Caucasian | 12.4 | NR | Anencephaly | NR | IIAM | | N | | | | | | • | • | • | • | • | | | | | | |
| DON65 | Neonatal | 1d | F | Black | 7.8 | NR | Anencephaly | NR | IIAM | | N | | | 0.967 | | | • | • | • | • | • | | | | | | |
| HDL066 | Neonatal | 1d | F | Unknown | NR | NR | Anencephaly | NR | | Pittsburgh | Y | | | | | | | | | | • | | | | | | |
| DON18 | Neonatal | 1d | F | Caucasian | 17.3 | NR | Exencephaly | NR | IIAM | | N | | | | | | • | • | • | • | • | | | | | | |
| HDL088 | Neonatal | 1d | F | Unknown | NR | NR | Anencephaly | 0 | OPO | | N | | | | | | | | | | • | | | | | | |
| HDL076 | Neonatal | 3d | F | Caucasian | 13.9 | NR | CVA/ICH | 1 | NDRI | | N | | | | | | | | | | • | | | | | | |
| HDL056 | Neonatal | 4d | M | Unknown | NR | NR | Anencephaly | NR | IIAM | | N | | | | | | | | | | • | | | | | | |
| HDL079 | Neonatal | 4d | M | Caucasian | NR | NR | Anoxia | NR | IIAM | | N | | | | | | | | | | • | | | | | | |
| DON22 | Neonatal | 4d | M | Caucasian | 13.1 | NR | Anoxia | NR | IIAM | | N | | | | | | • | • | • | • | • | | • | • | | | |
| DON8 | Neonatal | 5d | F | Caucasian | 16.5 | NR | Anencephaly | NR | IIAM | Pittsburgh | N | | | | | | • | • | • | • | • | • | | | | | |
| HDL034 | Neonatal | 8d | F | American Indian or Alaska Native | 10.9 | 4.4 | Anoxia | 4 | IIAM | | N | | | 0.330 | | | • | • | • | • | • | | | | • | | |
| HDL028 | Neonatal | 27d | F | Black | 20.0 | 5.2 | Anoxia | 6 | IIAM | | N | | | | | | • | • | • | • | • | | | | • | | |
| DON72 | Neonatal | 2m | M | Black | 13.4 | NR | Head trauma | 3 | IIAM | Pittsburgh | Y | | | | 14 | | • | • | • | • | • | • | • | | | • | • |
| HDL027 | Neonatal | 2m | F | Caucasian | 18.9 | NR | Anoxia | 1 | IIAM | | N | | | | | | • | • | • | • | | | | | | | |
| HDL061 | Neonatal | 2m | M | Black | 11.8 | NR | Anoxia | 4 | IIAM | | N | | | | | | | | | | • | | | | | | |
| HDL049 | Neonatal | 2m | F | Caucasian | 13.7 | 3.5 | Anoxia | 4 | IIAM | | N | | | | | | • | • | • | • | • | | | | • | | |
| DON29 | Neonatal | 2m | F | Hispanic or Latino | 20.8 | NR | Anoxia | NR | IIAM | Pittsburgh | N | | | | | | • | • | • | • | • | • | • | | | | |
| DON6 | Neonatal | 2m | F | Caucasian | 14.7 | NR | Head trauma | 3 | NDRI | | N | | | | | | • | • | • | • | • | • | • | | | | |
| HDL118 | Neonatal | 2m | M | Caucasian | 19.1 | 4.7 | Anoxia | 7 | IIAM | | N | | | | | | | | | | • | | | | | | |
| DON56 | Infancy | 3m | M | Caucasian | 20.5 | NR | Anoxia | 4 | NDRI | Pittsburgh | Y | | | | | | | | • | • | | • | • | | | • | • |
| DON3 | Infancy | 3m | M | Caucasian | 16.8 | NR | Anoxia | 14 | NDRI | | N | | | 0.250 | | | • | • | • | • | • | • | • | | | | |
| HDL055 | Infancy | 3m | M | Caucasian | 16.9 | 5.4 | Head trauma | 3 | OPO | | N | | | | | | • | • | • | • | • | | | | • | | |
| DON2 | Infancy | 3m | M | Hispanic or Latino | 16.1 | NR | Head trauma | 5 | NDRI | | N | | | | | | • | • | • | • | • | • | • | • | | | |
| HDL036 | Infancy | 3.75m | M | Hispanic or Latino | 13.0 | NR | Head trauma | NR | NDRI | | N | | | | | | • | • | • | • | • | | | | | | |
| HDL040 | Infancy | 4m | M | Caucasian | 18.8 | 5.5 | Anoxia | 3 | IIAM | | N | | | | | | • | • | • | • | • | | | | • | | |
| DON27 | Infancy | 7m | M | Hispanic or Latino | 17.3 | NR | Head trauma | NR | IIAM | | N | | | | | | • | • | • | • | | • | • | • | | | |
| HDL045 | Infancy | 8m | F | Hispanic or Latino | 21.1 | NR | Anoxia | 2 | IIAM | Pittsburgh | Y | | | | 10.37 | 10 | | | | | • | | | | | | |
| HDL104 | Infancy | 8m | F | Hispanic or Latino | 19.5 | 6.7 | CVA/ICH | 4 | nPOD | | N | | | | | | | | | | • | | | | | | |
| HDL082 | Infancy | 9m | F | Hispanic or Latino | 17.4 | 5.8 | Head trauma | 5 | IIAM | | N | | | | | | | | | | • | | | | | | |
| DON70 | Infancy | 10m | M | Black | 19.6 | NR | Anoxia | 2 | IIAM | Pittsburgh | Y | | | | | | • | • | • | • | | • | • | | | • | |
| DON60 | Infancy | 10m | F | Hispanic or Latino | 23.1 | NR | Head trauma | 3 | IIAM | Pittsburgh | Y | | | | | | • | • | • | • | • | • | • | | | • | • |
| DON10 | Infancy | 10m | F | Caucasian | 15.4 | NR | CVA/ICH | 4 | NDRI | | N | | | | | | • | • | • | • | • | • | • | | | | |
| DON31 | Infancy | 11m | M | Hispanic or Latino | 18.4 | 5.4 | Head trauma | 3 | NDRI | Pittsburgh | N | | | | | | • | • | • | • | • | • | • | | | | |
| HDL109 | Infancy | 13m | F | Caucasian | 18.4 | 5.1 | Anoxia | 3 | IIAM | | N | | | | | | | | | | • | | | | | | |
| HDL039 | Infancy | 13m | M | Hispanic or Latino | 25.2 | 5.6 | Anoxia | 6 | IIAM | | N | | | | | | • | • | • | • | • | | | | | | |
| HDL067 | Infancy | 13m | F | Caucasian | 18.8 | 5.2 | Head trauma | 4 | IIAM | | N | | | | | | | | | | • | | | | | | |
| HDL041 | Infancy | 15m | F | Caucasian | 16.7 | NR | Anoxia | 0 | IIAM | | N | | | | | | • | • | • | • | • | | | | • | | |
| HDL026 | Infancy | 15m | M | Black | 17.8 | 5.8 | Head trauma | 4 | IIAM | | N | | | | | | • | • | • | • | • | | | | | | |
| DON546 | Infancy | 16m | M | Caucasian | 24.4 | 5.1 | Head trauma | 0 | OPO | Imagine Pharma | Y | | | | | | | | | | | | | | | • | • |
| DON92 | Infancy | 16m | M | Caucasian | 22.0 | NR | Head trauma | 5 | NDRI | Pittsburgh | Y | | | | 11.75 | 64 | | | | | | | | | | • | • |
| DON534 | Infancy | 17m | M | Caucasian | 20.8 | NR | head trauma | 2 | OPO | Imagine Pharma | Y | | | | | | | | | | | | | | | • | |
| HDL004 | Infancy | 18m | M | Caucasian | 15.1 | 5.8 | Anoxia | 4 | OPO | | N | | | | | | • | • | • | • | • | | | | | | |
| HDL077 | Infancy | 18m | M | Hispanic or Latino | 15.6 | 5.9 | Anoxia | 6 | OPO | | N | | | | | | | | | | • | | | | | | |
| HDL086 | Infancy | 18m | F | Caucasian | 17.5 | 5.0 | Head trauma | 4 | IIAM | | N | | | | | | | | | | • | | | | | | |
| HDL042 | Infancy | 19m | F | Black | 16.8 | 5.7 | Head trauma | 3 | IIAM | Pittsburgh | Y | | | | 9.6 | 24 | | | | | | | | | | • | • |
| HDL070 | Infancy | 19m | F | Caucasian | 17.6 | NR | Anoxia | 0 | OPO | | N | | | 0.36 | | | | | | | • | | | | | | |
| HDL031 | Infancy | 19m | M | Caucasian | 16.5 | NR | Anoxia | 1 | NDRI | | N | | | | | | • | • | • | • | • | | | | | | |
| DON14 | Infancy | 19m | F | Hispanic or Latino | 12.9 | NR | Head trauma | 3 | NDRI | | N | | | | | | • | • | • | • | • | • | | | | | |
| DON12 | Infancy | 20m | F | Caucasian | 23.5 | NR | Anoxia | NR | IIAM | | N | | | | | | • | • | • | • | • | • | • | • | | | |
| HDL038 | Infancy | 20m | M | Hispanic or Latino | 14.8 | 5.7 | CVA/ICH | NR | NDRI | | N | | | | | | • | • | • | • | • | | | | | | |
| HDL063 | Infancy | 21m | F | Caucasian | 19.3 | 5.1 | Anoxia | 3 | IIAM | | N | | | | | | | | | | • | | | | | | |
| HDL009 | Infancy | 21m | M | Caucasian | 17.42 | 5.6 | Anoxia | 4 | IIAM | | N | | | | | | • | • | • | • | • | | | | | | |
| HDL008 | Infancy | 21m | M | Asian | 14.7 | NR | Anoxia | 7 | OPO | | N | | | | | | • | • | • | • | • | | | | • | | |
| DON28 | Infancy | 21m | M | Caucasian | 18.8 | NR | Anoxia | NR | IIAM | Pittsburgh | N | | | 0.450 | | | • | • | • | • | • | • | • | | | | |
| HDL158 | Childhood | 2y | M | Caucasian | 14.3 | 5.2 | Anoxia | 3 | IIAM | Imagine Pharma | Y | | | | | | | | | | | | | | | • | • |
| HDL129 | Childhood | 2y | M | Caucasian | 23.0 | 5.4 | Anoxia | 5 | IIAM | Imagine Pharma | Y | | | | | | | | | | | | | | | • | • |
| HDL053 | Childhood | 2y | M | Caucasian | 20.2 | NR | Anoxia | 3 | OPO | | N | | | | | | • | • | • | • | • | | | | | | |
| HDL064 | Childhood | 2y | M | Caucasian | 19.7 | 5.0 | Anoxia | 3 | IIAM | | N | | | | | | | | | | • | | | | | | |
| HDL035 | Childhood | 2y | M | Caucasian | 21.4 | 4.9 | Anoxia | 4 | IIAM | | N | | | | | | • | • | • | • | • | | | | | | |
| HDL030 | Childhood | 2y | M | Hispanic or Latino | 15.1 | 4.9 | Anoxia | 5 | OPO | | N | | | | | | • | • | • | • | • | | | | | | |
| HDL024 | Childhood | 2y | M | Hispanic or Latino | 14.4 | 5.7 | Anoxia | NR | IIAM | | N | | | | | | • | • | • | • | • | | | | • | | |
| HDL043 | Childhood | 2y | M | Caucasian | 15.7 | 5.4 | Head trauma | 1 | IIAM | | N | | | 0.4 | | | • | • | • | • | • | | | | | | |
| HDL015 | Childhood | 2y | F | Caucasian | 17.4 | 5.2 | Head trauma | 4 | OPO | | N | | | | | | • | • | • | • | • | | | | | | |
| HDL025 | Childhood | 2y | F | Caucasian | 16.9 | NR | Head trauma | 6 | IIAM | | N | | | | | | • | • | • | • | • | | | | | | |
| HDL033 | Childhood | 2y | F | Black | 17.8 | 4.7 | Head trauma | 10 | IIAM | | N | | | | | | • | • | • | • | • | | | | | | |
| HDL115 | Childhood | 2y | F | Caucasian | 18.5 | 5.2 | Anoxia | 4 | IIAM | | N | | | | | | | | | | • | | | | | | |
| HDL112 | Childhood | 3y | F | Caucasian | 17.8 | 5.1 | Head trauma | 3 | IIAM | | N | | | | | | | | | | • | | | | | | |
| HDL001 | Childhood | 3y | F | Caucasian | 12.0 | 6.0 | Head trauma | 2 | IIAM | Pittsburgh | Y | | | | 12 | 19 | • | • | • | • | • | | | | | • | • |
| HDL060 | Childhood | 3y | M | Caucasian | 16.8 | NR | Head trauma | 4 | IIAM | Pittsburgh | Y | | | | | | | | | | | | | | | • | |

**Table S1. Donor characteristics, sample types, and experimental usage**

Adapted from Hart NJ, Powers AC (2018), Checklist for Reporting Human Islet Preparations Used in Research, *Diabetologia*, doi.org/10.1007/s00125-018-4772-2.

| | Donor information | | | | | | | | | | | | | | | | Assays | | | | | | | | | | |
|---|---|---|---|---|---|---|---|---|---|---|---|---|---|---|---|---|---|---|---|---|---|---|---|---|---|---|---|
| Unique identifier | Age category | Age | Sex | Race/ethnicity | BMI (kg/m$^2$) | HbA1c (ng/mL) | Cause of death | Ventilator time (days) | Source of islets/ tissue | Islet isolation center (if applicable) | Function assessed? | Est. purity (%) | Hand-picked? | Warm ischemia time (hrs) | Cold ischemia time (hrs) | Culture time (hrs) | Endo dens | α/β/δ comp | α/β/δ prolif | α/β/δ dens | Panc weight | Innervation | Vasculature | Architecture | ε/γ comp | Perifusion | Islet gels |
| HDL012 | Childhood | 3y | F | Caucasian | 12.9 | 5.8 | Anoxia | 3 | OPO | Pittsburgh | N | | | | 14.033 | 17 | • | • | • | • | • | | | | • | | |
| DON46 | Childhood | 3y | M | Caucasian | 19.2 | NR | Anoxia | 5 | IIAM | Pittsburgh | N | | | | | | • | • | • | • | • | • | • | | | | |
| HDL014 | Childhood | 3y | F | Caucasian | 20.1 | NR | Head trauma | 1 | OPO | | N | | | | | | • | • | • | • | • | | | | | | |
| HDL013 | Childhood | 3y | F | Hispanic or Latino | 13.0 | 5.3 | Head trauma | 2 | NDRI | Pittsburgh | N | | | | | | • | • | • | • | • | | | | | | |
| HDL005 | Childhood | 4y | M | Caucasian | 19.7 | NR | Head trauma | NR | OPO | Pittsburgh | Y | | | | 7.48 | 4 | • | • | • | • | • | | | | | • | • |
| DON474 | Childhood | 4y | M | Caucasian | 15.39 | 5.5 | Head trauma | 2 | IIAM | Imagine Pharma | Y | | | | | | | | | | | | | | | • | • |
| DON13 | Childhood | 4y | F | Caucasian | 19.1 | NR | Head trauma | NR | IIAM | Pittsburgh | N | | | | | | • | • | • | • | • | • | • | • | | | |
| HDL029 | Childhood | 4y | F | Caucasian | 17.2 | 5.4 | Head trauma | NR | OPO | | N | | | | | | • | • | • | • | • | | | | | | |
| HDL126 | Childhood | 5y | F | Asian | 13.2 | 5.1 | Head trauma | 2 | IIAM | Imagine Pharma | Y | | | | 14.5 | | | | | | | | | | | • | • |
| DON7 | Childhood | 5y | M | Caucasian | 16.2 | 4.5 | Anoxia | 2 | IIAM | | N | | | | | | • | • | • | • | • | • | • | • | | | |
| DON1 | Childhood | 5y | F | Black | 15.9 | NR | Anoxia | NR | NDRI | | N | | | | | | • | • | • | • | • | • | • | | | | |
| DON50 | Childhood | 5y | M | Caucasian | 17.5 | NR | Head trauma | 3 | IIAM | Pittsburgh | N | | | | | | • | • | • | • | • | | | | | | |
| HDL016 | Childhood | 5y | M | Caucasian | 21.3 | 5.3 | Head trauma | 20 | OPO | | N | | | 0.92 | | | • | • | • | • | • | | | | | | |
| HDL084 | Childhood | 6y | F | Caucasian | 14.8 | NR | Anoxia | NR | OPO | Pittsburgh | Y | | | | 14.75 | 6 | | | | | • | | | | | • | • |
| HDL023 | Childhood | 6y | F | Hispanic or Latino | 18.2 | 6.2 | Anoxia | 10 | OPO | | N | | | | | | • | • | • | • | • | | | | • | | |
| HDL002 | Childhood | 6y | F | Black | 12.1 | NR | CVA/ICH | 15 | IIAM | | N | | | | | | • | • | • | • | • | | | | | | |
| DON43 | Childhood | 7y | M | Caucasian | 26.6 | NR | Anoxia | 2 | NDRI | Pittsburgh | Y | | | | | | • | • | • | • | • | • | • | | | • | • |
| HDL020 | Childhood | 7y | F | Hispanic or Latino | 15.0 | 5.2 | CVA/ICH | 2 | IIAM | Pittsburgh | Y | | | | 11.33 | 11 | • | • | • | • | • | | | | | • | |
| HDL017 | Childhood | 7y | F | Black | 20.8 | NR | Head trauma | 4 | NDRI | Pittsburgh | Y | | | | 15.767 | 1 | | | | | | | | | | • | |
| HDL051 | Childhood | 7y | F | Black | 18.4 | NR | Anoxia | NR | OPO | | N | | | | | | • | • | • | • | • | | | | | | |
| HDL019 | Childhood | 7y | M | Caucasian | 22.0 | 5.6 | Head trauma | NR | IIAM | | N | | | | | | • | • | • | • | • | | | | | | |
| DON91 | Childhood | 8y | M | Black | 17.2 | NR | Anoxia | 13 | NDRI | Pittsburgh | Y | | | | 12 | 16 | | | | | | | | | | • | |
| DON41 | Childhood | 8y | F | Caucasian | 16.1 | NR | CVA/ICH | NR | IIAM | Pittsburgh | Y | | | | | | • | • | • | • | • | | | | | • | |
| HDL083 | Childhood | 8y | M | Caucasian | 15.1 | 5.3 | Head trauma | 2 | IIAM | Pittsburgh | Y | | | | 12.28 | 6 | | | | | • | | | | | • | • |
| HDL032 | Childhood | 8y | F | Caucasian | 15.8 | 5.3 | Head trauma | NR | IIAM | Pittsburgh | Y | | | | 15.167 | 24 | • | • | • | • | • | | | | | • | • |
| HDL121 | Childhood | 8y | F | Black | 23.9 | NR | Head trauma | 2 | IIAM | Imagine Pharma | Y | | | | | | | | | | | | | | | • | • |
| HDL068 | Childhood | 8y | M | Caucasian | 15.7 | 5.3 | Anoxia | NR | OPO | Pittsburgh | N | | | | 15.88 | 10 | | | | | • | | | | | | |
| HDL021 | Childhood | 8y | M | Hispanic or Latino | 16.0 | 5.8 | Head trauma | 1 | OPO | | N | | | | | | • | • | • | • | • | | | | | | |
| HDL090 | Childhood | 8y | M | Black | 17.7 | 5.8 | Head trauma | 5 | IIAM | | N | | | | | | | | | | • | | | | | | |
| HDL052 | Childhood | 9y | M | Caucasian | 13.3 | 4.5 | Anoxia | 5 | IIAM | | N | | | | | | • | • | • | • | | | | | • | | |
| HDL006 | Childhood | 10y | M | Hispanic or Latino | 19.9 | 5.8 | Anoxia | 7 | OPO | Pittsburgh | Y | | | | | | • | • | | • | • | | | | | • | • |
| HDL010 | Childhood | 10y | F | Caucasian | 18.6 | 5.6 | Head trauma | 13 | OPO | Pittsburgh | Y | | | | 12.4 | 8 | • | • | • | • | • | | | | | • | • |
| HDL018 | Childhood | 10y | F | Hispanic or Latino | 25.4 | NR | Head trauma | NR | OPO | Pittsburgh | Y | | | | 14.62 | 20 | | | | | | | | | | • | • |
| DON142 | Childhood | 10y | M | Caucasian | 23.8 | 6.0 | Head trauma | NR | IIAM | Pittsburgh | Y | | | | 20.85 | 44 | | | | | | | | | | • | |
| DON4 | Childhood | 10y | M | Caucasian | 19.3 | NR | Head trauma | 7 | NDRI | | N | | | | | | • | • | • | • | • | • | • | • | | | |
| DON328 | Adult | 24y | M | Caucasian | 31.7 | NR | Anoxia | 1 | OPO | Pittsburgh | Y | | | 0.28 | 5.92 | 24 | | | | | | | | | | • | |
| DON382 | Adult | 27y | M | Caucasian | 40.6 | NR | Head trauma | NR | IIAM | Pittsburgh | Y | | | | 12 | 3.5 | | | | | | | | | | • | |
| DON481 | Adult | 28y | M | Caucasian | 25.1 | 5.5 | Head trauma | 4 | IIAM | Imagine Pharma | Y | | | | | | | | | | | | | | | • | |
| DON420 | Adult | 30y | M | Caucasian | 25.9 | 5.4 | Head trauma | 3 | IIAM | Pittsburgh | Y | | | | | | | | | | | | | | | • | |
| DON308 | Adult | 35y | M | Caucasian | 24.7 | 5.3 | Head trauma | 6 | OPO | Pittsburgh | Y | | | | 6.18 | 19 | | | | | | | | | | • | |
| DON246 | Adult | 42y | M | Caucasian | 32.2 | 6.0 | Overdose | NR | OPO | Pittsburgh | Y | | | | 14.5 | 10 | | | | | | | | | | • | |
| DON227 | Adult | 45y | F | Caucasian | 29.8 | 5.6 | Anoxia | 3 | OPO | Pittsburgh | Y | | | | 9 | 23 | | | | | | | | | | • | |
| DON389 | Adult | 46y | F | Caucasian | 32.9 | 5.7 | CVA/ICH | NR | IIAM | Pittsburgh | Y | | | | 8.2 | 48 | | | | | | | | | | • | |
| DON316 | Adult | 48y | M | Caucasian | 24.6 | 4.9 | Anoxia | 4 | OPO | Pittsburgh | Y | | | | 20.48 | 27.5 | | | | | | | | | | • | |
| DON204 | Adult | 52y | M | Black | 29.2 | NR | CVA/ICH | NR | OPO | Pittsburgh | Y | | | | 8.7 | 23 | | | | | | | | | | • | |
| DON486 | Adult | 55y | M | Caucasian | 35.5 | 4.9 | Anoxia | 5 | IIAM | Imagine Pharma | Y | | | | 44.27 | | | | | | | | | | | • | |
| DON61 | Adult | 55y | M | Black | 35.6 | NR | CVA/ICH | 4 | IIAM | Pittsburgh | Y | | | | | | | | | | | | | | | • | |
| DON197 | Adult | 55y | F | Black | 24.2 | NR | CVA/ICH | 1 | OPO | Pittsburgh | Y | | | 0 | 3.8 | 28 | | | | | | | | | | • | |
| DON381 | Adult | 59y | M | Caucasian | 32.7 | 5.5 | Head trauma | 1 | IIAM | Pittsburgh | Y | | | | | | | | | | | | | | | • | |

*Abbreviations: CVA, cardiovascular accident; d, days; F, female; G, gestational age; ICH, intracerebral hemorrhage; IHC, immunohistochemistry; IIAM, International Institute for the Advancement of Medicine; IIDP, Integrated Islet Distribution Program; m, months; M, male; N/A, not available; NDRI, National Disease Research Interchange; OPO, organ procurement organization (local); Perif., perifusion; w, weeks.*
