## Supplemental Table S2 for "Mapping histologic and functional maturation of human endocrine pancreas across early postnatal periods"

**Table S2. Primary antibodies used for immunohistochemistry**

| Antigen/Conjugate | Species | Source | Catalog # | RRID | IF | ABA | TSA | Whole mount |
|---|---|---|---|---|---|---|---|---|
| C-peptide | Rat | DSHB | GN-ID4 | AB_2255626 | 1:100 | | | |
| CD31 (PECAM1) | Mouse | BD Biosciences | 550389 | AB_2252087 | 1:100 (1:50) | | | 1:50 |
| Collagen IV | Rabbit | Abcam | ab6586 | AB_305584 | 1:1000 | | | |
| Ghrelin | Mouse | Abcam | ab57222 | AB_941760 | 1:5000 | | | |
| Ghrelin | Rat | R&D Systems | MAB8200 | AB_2637039 | 1:1000 | | | |
| Glucagon | Mouse | Abcam | ab10988 | AB_297642 | 1:500 | | | |
| Glucagon | Rabbit | Cell Signaling | 2760s | AB_659831 | 1:10k | | | 1:100 |
| Glucagon | Guinea pig | Linco (Millipore) | 4031 | AB_433707 | 1:1k-10k | | | 1:10000 |
| Glucagon | Rabbit | Linco (Millipore) | 4030 | AB_1977523 | 1:1000 | | | |
| GP2 | Mouse | MBL International | D277-3 | AB_10598500 | 1:250 | | | |
| GP2 (Hpx1) | Mouse | P. Streeter | N/A | | 1:100 | | | |
| Insulin | Guinea pig | Abcam | ab7842 | AB_306130 | 1:1000 | | | |
| Insulin | Guinea pig | Cell Marque | 273A-14 | AB_1158520 | 1:100-500 | | | |
| Insulin | Guinea pig | Dako (Agilent) | A0564 | AB_10013624 | 1:1000 | | | |
| Insulin | Guinea pig | Linco (Millipore) | 4010 | AB_1587205 | 1:1000 | | | 1:1000 |
| Insulin | Goat | Santa Cruz | sc7839 | AB_2296108 | 1:100 | | | |
| Ki67 | Rabbit | Abcam | ab15580 | AB_443209 | 1:5000 | | | |
| Nkx6.1 | Rabbit | BCBC (P. Serup) | AB1069 | | 1:1000-2000 | | | |
| Nkx6.1 | Mouse | DSHB | F55A12 | AB_532379 | | 1:100 | 1:500 | |
| Pancreatic polypeptide | Rabbit | Bachem | T-4088 | AB_518533 | 1:1000 | | | |
| Pancreatic polypeptide | Mouse | R&D Systems | MAB62971 | AB_11127208 | 1:100 | | | |
| Pax6 | Rabbit | BioLegend | 901302 | AB_2565003 | 1:2000 | | | |
| Pax6 | Rabbit | Covance | PRB-278P | AB_291612 | 1:200 | 1:500 | 1:1000 | |
| Ptf1a | Rabbit | BCBC | N/A | | | | 1:1000 | |
| Ptf1a | Goat | C. Wright? | N/A | | 1:500 | | | |
| Somatostatin | Sheep | American Research Products | 13-2366 | AB_1542966 | 1:500 | | | 1:500 |
| Somatostatin | Goat | Santa Cruz | sc7819 | AB_2302603 | 1:500-1000 | | | |
| Synapsin I/II | Rabbit | Synaptic Systems | 106002 | AB_887804 | 1:2000 | | | |
| Tuj1 (TUBB3) | Rabbit | Covance | MRB-435P | AB_663339 | 1:20k | | | 1:20000 |

ABA, avidin-biotin amplification; IF, immunofluorescence; TSA, tyramide signal amplification. All antibodies were used on 8-10 μm cryosections; if dilution factor differed for thick sections (30 μm) and/or whole-mounts, a second dilution factor is listed in parentheses.
