## Supplemental Table S3 for "Mapping histologic and functional maturation of human endocrine pancreas across early postnatal periods"

**Table S3. Secondary antibodies used for immunohistochemistry**

| Target species/isotype | Host species | Conjugate | Source | Catalog # | IF |
|---|---|---|---|---|---|
| Goat IgG | Donkey | Cy5 | Jackson Immunoresearch | 705-175-147 | 1:200 |
| Goat IgG | Donkey | Cy2 | Jackson Immunoresearch | 705-225-147 | 1:500 |
| Goat IgG | Donkey | Cy3 | Jackson Immunoresearch | 705-165-147 | 1:500 |
| Guinea pig IgG | Donkey | AF 488 | Jackson Immunoresearch | 706-545-148 | 1:500 |
| Guinea pig IgG | Donkey | AF 647 | Jackson Immunoresearch | 706-605-148 | 1:200 |
| Guinea pig IgG | Donkey | Cy2 | Jackson Immunoresearch | 706-225-148 | 1:500 |
| Guinea pig IgG | Donkey | Cy3 | Jackson Immunoresearch | 706-165-148 | 1:500 |
| Guinea pig IgG | Donkey | Cy5 | Jackson Immunoresearch | 706-175-148 | 1:200 |
| Mouse IgG | Donkey | Cy2 | Jackson Immunoresearch | 715-225-150 | 1:500 |
| Mouse IgG | Donkey | Cy3 | Jackson Immunoresearch | 715-165-150 | 1:500 |
| Mouse IgG | Donkey | Cy5 | Jackson Immunoresearch | 715-175-150 | 1:200 |
| Rabbit IgG | Donkey | AF 488 | Jackson Immunoresearch | 711-545-152 | 1:500 |
| Rabbit IgG | Donkey | Cy2 | Jackson Immunoresearch | 711-225-152 | 1:500 |
| Rabbit IgG | Donkey | Cy3 | Jackson Immunoresearch | 711-165-152 | 1:500 |
| Rabbit IgG | Donkey | Cy5 | Jackson Immunoresearch | 711-175-152 | 1:200 |
| Rat IgG | Donkey | AF 488 | Jackson Immunoresearch | 712-545-153 | 1:500 |
| Rat IgG | Donkey | Cy2 | Jackson Immunoresearch | 712-225-153 | 1:500 |
| Rat IgG | Donkey | Cy3 | Jackson Immunoresearch | 712-165-153 | 1:500 |
| Rat IgG | Donkey | Cy5 | Jackson Immunoresearch | 712-175-153 | 1:200 |
| Sheep IgG | Donkey | Cy2 | Jackson Immunoresearch | 713-225-147 | 1:500 |
| Sheep IgG | Donkey | Cy3 | Jackson Immunoresearch | 713-165-147 | 1:500 |
| Sheep IgG | Donkey | Cy5 | Jackson Immunoresearch | 713-175-147 | 1:200 |
