## Supplemental Table S4 for "Mapping histologic and functional maturation of human endocrine pancreas across early postnatal periods"

**Table S4. Hormone assays used for assessing islet function**

| | | | | Islet hormone content data inclusion | | Perifusion trace data inclusion | | | |
|---|---|---|---|---|---|---|---|---|---|
| **Donor ID** | **Group** | **INS assay(s)** | **GCG assay(s)** | **INS** | **GCG** | **INS (IEQ)** | **INS (cont)** | **GCG (IEQ)** | **GCG (cont)** |
| DON72 | Infancy | Millipore RIA | R&D ELISA, Mercodia ELISA | Included | No extract | Included | Included | Included | No extract |
| DON56 | Infancy | Millipore RIA | R&D ELISA, Mercodia ELISA | Included | Below detection (extract) | Included | Included | Included | Below detection (extract) |
| DON60 | Infancy | Millipore RIA | R&D ELISA, Mercodia ELISA | Included | Included | Included | Included | Included | Below detection (extract) |
| DON70 | Infancy | Millipore RIA | R&D ELISA | Included | Below detection (extract) | Included | Included | Below detection | Below detection (extract) |
| DON546 | Infancy | Mercodia ELISA | R&D ELISA, Mercodia ELISA | Included | Included | Included | Included | Included | Included |
| DON92 | Infancy | Millipore RIA | R&D ELISA, Mercodia ELISA | Included | No extract | Included | Included | Included | No extract |
| DON534 | Infancy | Mercodia ELISA | R&D ELISA | Included | Included | Included | Included | Included | Included |
| HDL042 | Infancy | Millipore RIA | R&D ELISA, Mercodia ELISA, Millipore RIA | Included | Included | Included | Included | Included | Included |
| HDL129 | Childhood | Mercodia ELISA | R&D ELISA | Included | Included | Included | Included | Included | Included |
| HDL158 | Childhood | Mercodia ELISA | R&D ELISA, Mercodia ELISA | Included | Included | Included | Included | Included | Included |
| HDL001 | Childhood | Millipore RIA | Millipore RIA | Included | Included | Included | Included | Included | Included |
| HDL060 | Childhood | Millipore RIA | Millipore RIA | Included | Included | Included | Included | Included | Included |
| HDL005 | Childhood | Millipore RIA | Millipore RIA | Included | Included | Included | Included | Included | Included |
| DON474 | Childhood | Mercodia ELISA | R&D ELISA | Included | Included | Included | Included | Included | Included |
| HDL126 | Childhood | Mercodia ELISA | R&D ELISA | Included | Included | Included | Included | Included | Included |
| HDL084 | Childhood | Millipore RIA | Millipore RIA | Included | Included | Included | Included | Included | Included |
| HDL020 | Childhood | Millipore RIA | Millipore RIA | Included | Included | Included | Included | Included | Included |
| HDL017 | Childhood | Millipore RIA | Millipore RIA | Included | Included | Included | Included | Included | Included |
| DON43 | Childhood | Millipore RIA | R&D ELISA | Included | Included | Included | Included | Included | Excluded (outlier) |
| HDL032 | Childhood | Millipore RIA | Millipore RIA | Included | Included | Included | Included | Included | Included |
| HDL083 | Childhood | Millipore RIA | Millipore RIA | Included | Included | Included | Included | Included | Included |
| HDL121 | Childhood | Mercodia ELISA | Cisbio HTRF | Included | Included | Included | Included | Included | Included |
| DON91 | Childhood | Millipore RIA | Millipore RIA | Included | Included | Included | Included | Included | Included |
| DON41 | Childhood | Millipore RIA | Millipore RIA | Included | Included | Included | Included | Included | Included |
| HDL006 | Childhood | Millipore RIA | Millipore RIA | Pellet loss | Pellet loss | Included | Pellet loss | Included | Pellet loss |
| HDL018 | Childhood | Millipore RIA | Millipore RIA | Included | Included | Included | Included | Included | Included |
| HDL010 | Childhood | Millipore RIA | Millipore RIA | Included | Included | Included | Included | Included | Included |
| DON142 | Childhood | Millipore RIA | R&D ELISA, Mercodia ELISA | Included | Included | Included | Included | Included | Included |
| DON61 | Adult | Mercodia RIA | Mercodia RIA | Included | Included | Included | Included | Included | Included |
| DON197 | Adult | Mercodia RIA | Mercodia RIA | Included | Included | Included | Included | Included | Included |
| DON204 | Adult | Mercodia RIA | Mercodia RIA | Included | Included | Included | Included | Included | Included |
| DON227 | Adult | Mercodia RIA | Mercodia RIA | Included | Included | Included | Included | Excluded (abnormal trace) | Included |
| DON246 | Adult | Mercodia RIA | Mercodia RIA | Included | Included | Included | Included | Included | Included |
| DON308 | Adult | Mercodia RIA | Mercodia RIA | Included | Included | Included | Included | Included | Included |
| DON316 | Adult | Mercodia RIA | Mercodia RIA | Included | Included | Included | Included | Included | Included |
| DON328 | Adult | Mercodia RIA | Mercodia RIA | Included | No extract | Included | Included | Included | No extract |
| DON382 | Adult | Mercodia RIA | Mercodia RIA | Included | Included | Excluded (abnormal trace) | Included | Excluded (low response) | Excluded (low response) |
| DON381 | Adult | Mercodia RIA | Cisbio HTRF | Included | Included | Included | Included | Included | Included |
| DON389 | Adult | Mercodia ELISA | Cisbio HTRF | Included | Included | Included | Included | Included | Included |
| DON420 | Adult | Mercodia ELISA | Cisbio HTRF | Included | Included | Excluded (low response) | Included | Included | Included |
| DON481 | Adult | Mercodia ELISA | R&D ELISA | Included | Included | Included | Included | Excluded (abnormal trace) | Included |
| DON486 | Adult | Mercodia ELISA | R&D ELISA | Included | Included | Included | Included | Included | Included |

**Human insulin assays:** Millipore RIA RI-13K, Mercodia ELISA 10-1132-01

**Glucagon assays:** Millipore RIA GL-32K, Mercodia ELISA 10-1281-01, R&D ELISA DGCG0, Cisbio HTRF 62CGLPEH
