## Supplemental Table S5 for "Mapping histologic and functional maturation of human endocrine pancreas across early postnatal periods"

**Table S5. Comparison of dynamic perifusion profile between infancy, childhood, and adult samples**

| Hormone detected | Normalization | Group 1 | Group 2 | L ratio | p-value |
|---|---|---|---|---|---|
| Insulin | IEQ | Global | - | 219.0366 | 0 |
| Insulin | IEQ | Infancy | Childhood | 150.5594 | 0 |
| Insulin | IEQ | Infancy | Adult | 114.6687 | 0 |
| Insulin | IEQ | Childhood | Adult | 173.0602 | 0 |
| Glucagon | IEQ | Global | - | 146.9903 | 0 |
| Glucagon | IEQ | Infancy | Childhood | 92.0964 | 0 |
| Glucagon | IEQ | Infancy | Adult | 86.6722 | 0 |
| Glucagon | IEQ | Childhood | Adult | 98.9603 | 0 |
| Insulin | Insulin content | Global | - | 109.7987 | 0 |
| Insulin | Insulin content | Infancy | Childhood | 75.2847 | 0 |
| Insulin | Insulin content | Infancy | Adult | 63.0174 | 0 |
| Insulin | Insulin content | Childhood | Adult | 83.7083 | 0 |
| Glucagon | Glucagon content | Global | - | 51.0292 | 0 |
| Glucagon | Glucagon content | Infancy | Childhood | 27.9202 | 0.0019 |
| Glucagon | Glucagon content | Infancy | Adult | 33.7995 | 0.0002 |
| Glucagon | Glucagon content | Childhood | Adult | 29.4132 | 0.0011 |
